# Metformin Enhances Antibody-Mediated Recognition of HIV-Infected CD4^+^ T-Cells by Decreasing Viral Release

**DOI:** 10.1101/2024.02.15.580166

**Authors:** Augustine Fert, Jonathan Richard, Laurence Raymond Marchand, Delphine Planas, Jean-Pierre Routy, Nicolas Chomont, Andrés Finzi, Petronela Ancuta

## Abstract

The mechanistic target of rapamycin (mTOR) positively regulates multiple steps of the HIV-1 replication cycle. We previously reported that a 12-weeks supplementation of antiretroviral therapy (ART) with metformin, an indirect mTOR inhibitor used in type-2 diabetes treatment, reduced mTOR activation and HIV transcription in colon-infiltrating CD4^+^ T-cells, together with systemic inflammation in nondiabetic people with HIV-1 (PWH). Herein, we investigated the antiviral mechanisms of metformin. In a viral outgrowth assay performed with CD4^+^ T-cells from ART-treated PWH, and upon infection *in vitro* with replication-competent and VSV-G-pseudotyped HIV-1, metformin decreased virion release, but increased the frequency of productively infected CD4^low^HIV-p24^+^ T-cells. These observations coincided with increased BST2/Tetherin (HIV release inhibitor) and Bcl-2 (pro-survival factor) expression, and improved recognition of productively infected T-cells by HIV-1 Envelope antibodies. Thus, metformin exerts pleiotropic effects on post-transcription/translation steps of the HIV-1 replication cycle and may be used to accelerate viral reservoir decay in ART-treated PWH.

**Graphical Abstract.**
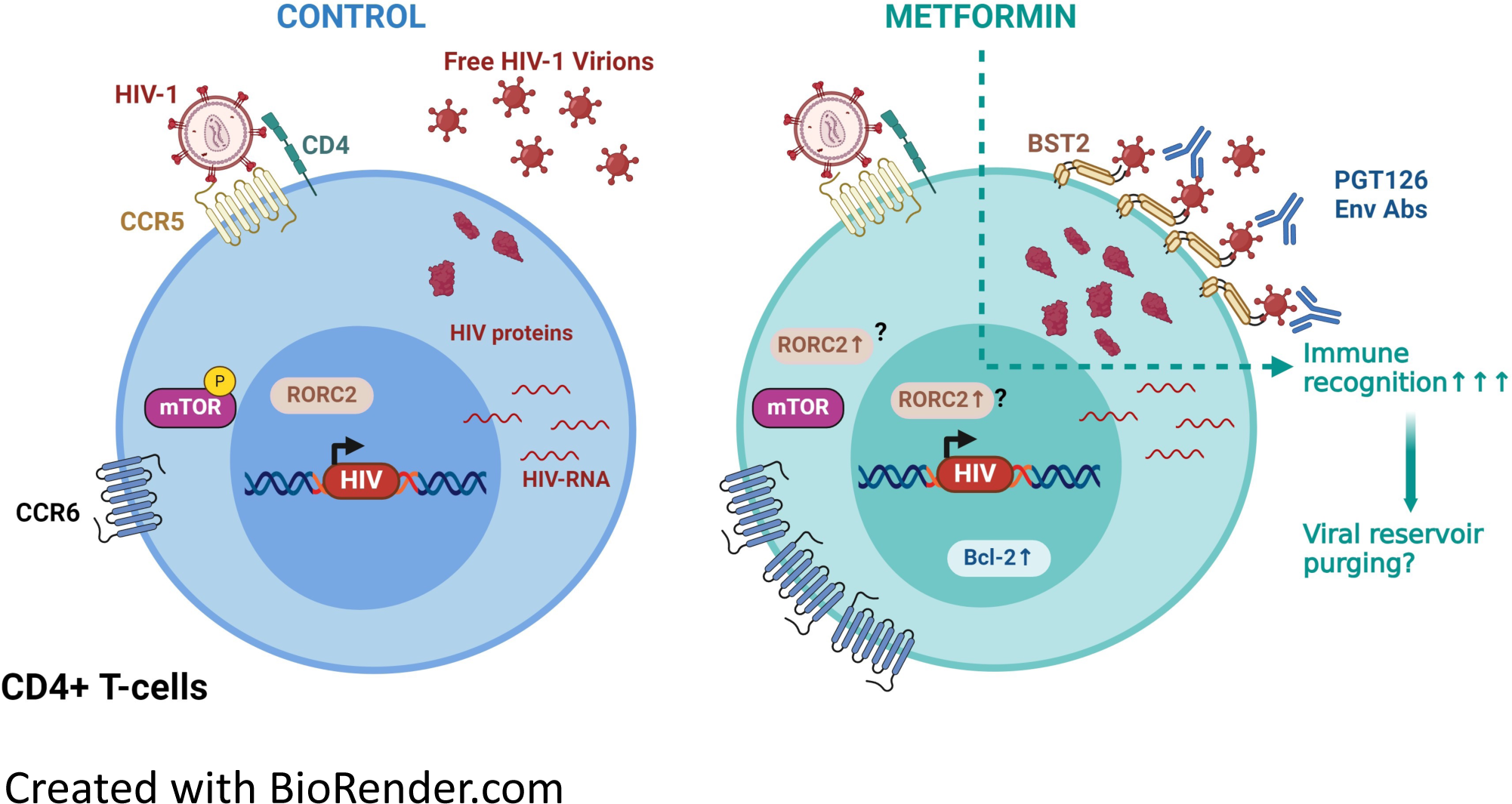

## INTRODUCTION

Antiretroviral therapy (ART) efficiently reduces HIV-1 replication to undetectable plasma levels and increases the life quality of people with HIV-1 (PWH)^1^. However, ART does not eradicate HIV-1 since viral reservoirs (VRs) persist in cells and tissues, a process associated with increased risk of developing non-AIDS comorbidities such as cardiovascular diseases, cancers and metabolic disorders, thus causing accelerated aging (*i.e.,* frailty, dementia)^2–8^. Those HIV-1 related pathologies are caused by chronic immune activation^9–11^. Chronic immune activation in PWH is multifactorial. In fact, suboptimal ART penetration may occur in specific tissues and cell subsets^12^. In addition, ART does not restore mucosal CD4^+^ T-cells populations depleted by HIV-1 infection^13^. Finally, ART does not block HIV-1 transcription and translation leading to residual HIV-RNA and protein expression^12,14,15^. Those byproducts participate to the chronic activation of the immune system^9,16^. New therapeutic strategies are needed to reduce comorbidities associated with chronic inflammation in PWH. In the absence of an HIV cure, whether strategies targeting HIV transcription and translation could achieve this goal remains to be determined.

The efficacy of HIV-1 replication depends on the biological features of target cells, including their metabolic state^17,18^. Indeed, subsets of CD4^+^ T-cells with increased metabolic activity, such as effector memory and Th17-polarized CCR6^+^ CD4^+^ T-cells, represent major targets for HIV-1 replication^17,19–22^. Glycolysis and oxidative phosphorylation (OXPHOS) are two major metabolic pathways that provide energy to cells^23^. HIV-1 replication is an energy-intensive process, this requirement is met by increasing the glycolysis to produce energy^24^. Accordingly, transcriptional profiling of CD4^+^ T-cells from the RV217 study cohort showed that HIV-1 plasma viral load positively correlated with the expression of genes involved in the glycolysis and OXPHOS^25^. The fact that energy fueling metabolic activities in HIV-infected cells depend more on glycolysis, compared to the metabolic status of uninfected cells, suggests the potential use of immunometabolism modulators (glycolysis/OXPHOS inhibitors) as new therapeutic interventions to limit the production of viral byproducts by stable viral reservoirs in ART-treated PWH^7,26,27^.

The mechanistic Target Of Rapamycin (mTOR) pathway, a key regulator of T-cell differentiation and growth *via* the induction of glycolysis^28–30^, was reported to be involved in multiple steps of the HIV-1 replication cycle (*i.e.,* entry, reverse transcription, nuclear transport and transcription)^31–34^. Previous work by our group demonstrated that mTOR expression and phosphorylation was preferentially induced in CD4^+^ T-cells expressing the Th17 marker CCR6 upon T-cell Receptor (TCR) triggering and exposure to the gut-homing modulator retinoic acid (RA) *via* mechanisms involving the upregulation of the HIV-1 co-receptor CCR5^20^. Also, by using a VSV-G-pseudotyped HIV-1 construct, which enters cells *via* endocytosis independently of CD4 and co-receptors^35^, we demonstrated that mTOR activation facilitates the post-entry steps of the HIV-1 replication cycle in RA-treated CCR6^+^CD4^+^ T-cells^20^. Furthermore, we demonstrated that mTOR inhibitors reduce viral outgrowth in CCR6^+^CD4^+^ T-cells of ART-treated PWH^20^. Consistently, Besnard *et al.,* showed that mTOR activation promotes HIV-1 transcription, *via* mechanisms involving the phosphorylating of CDK9, a subunit of the PTFEb complex, needed for HIV-1 transcription elongation^33^. Finally, Taylor *et al.,* showed that mTOR activity is increased in CD4^+^ T-cells from ART-PWH compared to HIV-uninfected individuals; mTOR inhibition in TCR-activated CD4^+^ T-cells leads to a decrease in the pool of dNTPs needed for HIV-1 reverse transcription; and that mTOR activation stabilizes microtubules in HIV-infected T-cells to facilitate the nuclear import of HIV-1 pre-integration complexes^34^.

Knowledge on the importance of mTOR pathway in regulating specific steps of the HIV-1 replication cycle^20,31–34^, prompted mTOR targeting *in vivo* in ART-treated PWH. In a single-arm clinical trial, 6-month supplementation of ART with everolimus, a direct mTOR inhibitor used as immunosuppressive drug in transplant recipient^36^, decreased mTOR activation, as well as the cell-associated (CA) HIV-RNA levels in blood CD4^+^ T-cells^37^, thus pointing to a potential direct role of mTOR in modulating HIV transcription *in vivo*. However, the use of such immunosuppressive drugs is not recommended outside organ transplantation. Metformin, an indirect mTOR inhibitor, is a drug approved by the Food and Drug Administration (FDA) and widely used to treat type-2 diabetes, as well as other metabolic disorders^38,39^. The repurposing of metformin for cancer is currently studied^40,41^. Mechanistically, metformin blocks the first complex of the respiratory chain of the mitochondria, leading to an increase in the AMP/ATP ratio. The change in this ratio leads to an activation of AMP-activated Protein Kinase (AMPK) pathway, which results in mTOR pathway inhibition^42–44^. Recently, our group performed a pilot non-randomized clinical trial in which non-diabetic ART-treated PWH received metformin for 12 weeks, and matched blood and colon biopsies were collected at baseline and the end of treatment for immunological and virological measurements^45,46^. Our results revealed that mTOR was preferentially expressed in CCR6^+^CD4^+^ T-cells from the colon of ART-treated PWH at baseline. Moreover, metformin significantly decreased the frequency of colon-infiltrating CD4^+^ T-cells and mTOR phosphorylation in CCR6^+^CD4^+^ T-cells, and reduced levels of systemic inflammation (*i.e.,* sCD14). Finally, a reduction in HIV-1 transcription, measured as the HIV-RNA/DNA ratio, was observed in CD4^+^ T-cells isolated from the colon in a fraction of study participants (8/13), thus pointing to a potential link between mTOR activation and HIV-1 transcription in T-cells carrying VRs^46^. Of interest, Guo *et al.,* showed that metformin reduced HIV-1 replication and limited CD4^+^ T-cells depletion in a humanized mouse model reconstituted with primary human CD4^+^ T-cells^25^. Another study showed that 24 weeks of metformin administration in complement of ART decreased the frequency of CD4^+^ T-cells with a PD1^+^TIGIT^+^TIM3^+^ phenotype considered as a molecular signature for exhausted cells contributing to HIV persistence^47–49^. Whether the metformin supplementation of ART may represent a valuable strategy to decrease immune dysfunction in PWH requires further investigations.

Given the encouraging results of metformin supplementation of ART in PWH^46,50^, we sought to identify the steps of the HIV-1 replication cycle modulated by metformin. To this aim, we studied the effects of metformin in a viral outgrowth assay (VOA) monitored in CD4^+^ T-cells of ART-treated PWH, and performed infection of CD4^+^ T-cells from HIV-uninfected participants *in vitro* using replication-competent and single-round VSV-G-pseudotyped HIV-1. Our results demonstrate that metformin exerts antiviral effects by blocking the release of HIV-1 virions *in vitro*, despite an unexpected capacity to increase HIV-p24 expression at single-cell level. Similarly, metformin boosted HIV-1 outgrowth without increasing viral release from reactivated reservoir cells, *via* mechanisms involving increased BST2 expression. Of note, metformin favored the recognition of reactivated reservoir cells by broadly neutralizing (bNAbs) anti-HIV-Env Abs. Overall, these results reveal the pleiotropic effects of metformin on post-integration steps of the viral replication cycle and support a model in which metformin may be used to accelerate viral reservoir decay during ART, especially in combination with immune interventions aimed at boosting anti-HIV immunity, such as the antibody-dependent cellular cytotoxicity (ADCC).

## RESULTS

### Metformin promotes VR reactivation in memory CD4^+^ T-cells of ART-treated PWH

To determine the optimal concentration of metformin for experiments *in vitro*, we first tested the effect of different doses of metformin on the phosphorylation of mTOR and its downstream substrate, the ribosomal S6 kinase (S6K). Results demonstrate that metformin at 1 mM efficiently inhibited mTOR and S6K phosphorylation upon TCR triggering, without impacting on cell viability and proliferation (Supplemental Figure 1). Metformin at 1 mM also showed efficacy in blocking the mTOR pathway in another study using CD4^+^ T-cells^25^. To explore the effect of metformin on HIV-1 reservoirs reactivation, we performed a simplified viral outgrowth assay (VOA), as depicted in Figure 1A^51^. INK128, a potent direct mTORC1/mTORC2 inhibitor^52^, was used in parallel.

**Figure 1:**
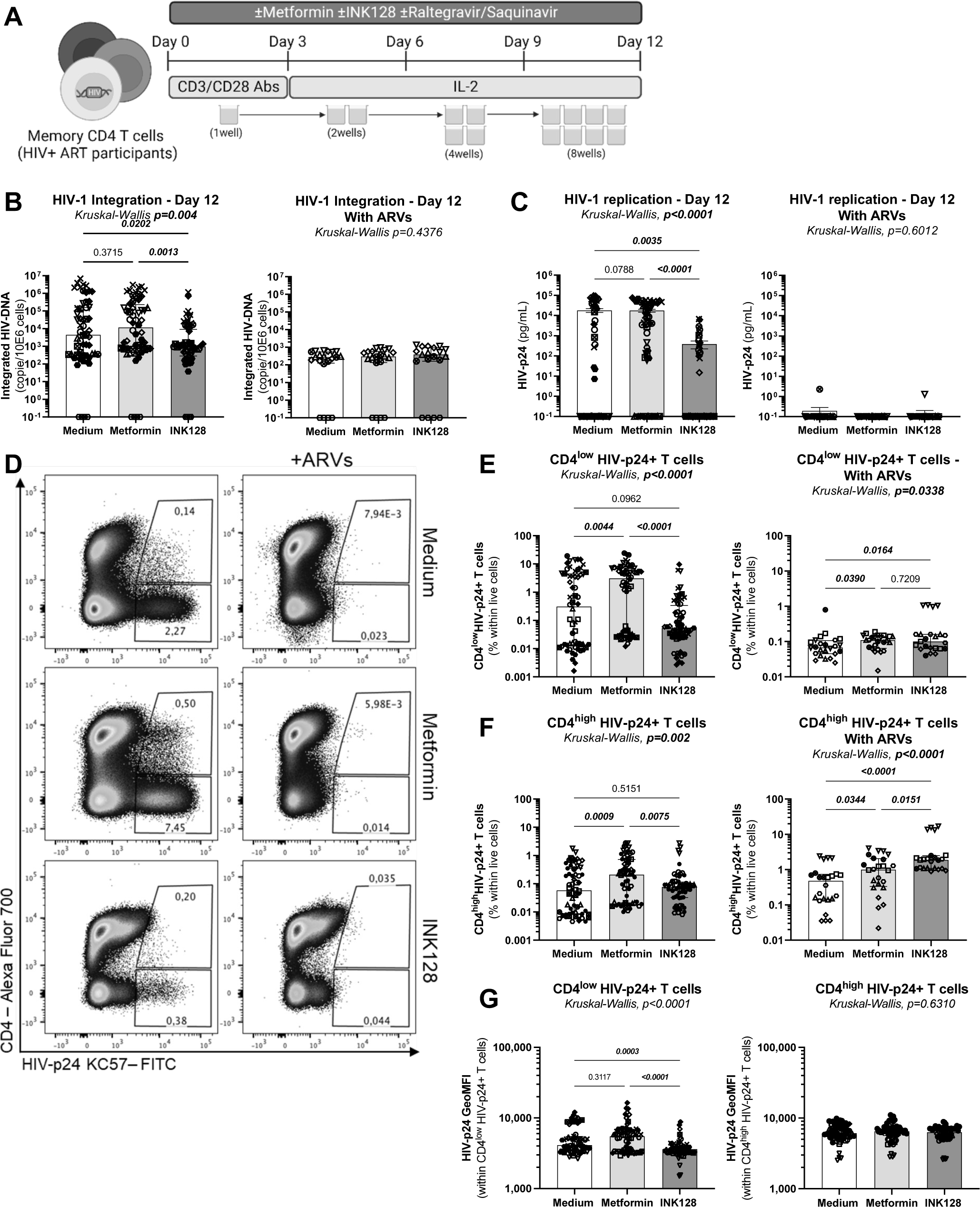
Effects of metformin on viral outgrowth in memory CD4^+^ T-cells from ART-treated PWH. **(A)** Shown is the flow chart for the viral outgrowth assay (VOA). Briefly, memory CD4^+^ T-cells from ART-treated PWH were stimulated with CD3/CD28 Abs, in the presence (n=11) or the absence (n=6) of ARVs and in the presence/absence of metformin (1 mM) or INK128 (50 nM) for 3 days. Supernatants were collected, cells were split in two news wells, and media containing IL-2 and metformin or INK128 was refreshed every 3 days. Experiments were performed in 4-8 original replicates *per* condition. One original replicate at day 0 generated 8 final replicates at day 12. At the end of the experiment, T-cells derived from the same original replicate were merged for PCR and flow cytometry analysis. Shown are **(B)** integrated HIV-DNA levels quantified by real-time nested PCR and **(C)** levels of HIV-p24 in cell-culture supernatants quantified by ELISA. Shown are **(D)** representative flow cytometry dot plot of intracellular HIV-p24 and surface CD4 expression; **(E)** statistical analysis of the frequency of CD4^low^HIV-p24^+^, **(F)** the frequency of CD4^high^HIV-p24^+^ cells; and **(G)** the geometric MFI (GeoMFI) of HIV-p24 expression in CD4^low^ HIV-p24^+^ and CD4^high^ HIV-p24^+^ T-cell subsets. Each symbol represents one donor; bars indicate the median ± interquartile range. Kruskal-Wallis test and uncorrected Dunn’s multiple comparison p-values are indicated on the graphs.

When the VOA was performed in the absence of ARVs using CD4^+^ T-cells from ART-treated PWH (n=11), the mTOR inhibitor INK128 reduced viral outgrowth, as reflected by levels of integrated HIV-DNA in cells and HIV-p24 levels in cell-culture supernatants (Figure 1B-C<u>, left panels</u>), consistent with the antiviral effects of INK128 previously reported by our group and others^20,31^. Unexpectedly, under these same experimental conditions, metformin did not affect viral outgrowth, as measured by PCR in cells and ELISA in cell-culture supernatants (Figure 1B-C<u>, left panels</u>). In contrast, metformin increased the frequency of productively infected cells, identified as cells with a CD4^low^HIV-p24^+^ phenotype (Figure 1D-E<u>, left panels</u>). Metformin also increased the frequency of a relatively small subset of CD4^high^HIV-p24^+^ T-cells, which may represent cells recently coated with HIV virions^53^ (Figure 1D and F<u>, left panels</u>). There was no increase in the geometric MFI of HIV-p24 expression within CD4^low^HIV-p24^+^ and CD4^high^HIV-p24^+^ T-cells (Figure 1G<u>, left panel</u>). These results demonstrate that metformin facilitates the expansion of productively infected cells upon TCR-triggering *in vitro*, without affecting the accumulation of HIV-p24 inside infected cells, nor the release of free progeny virions in cell-culture supernatants.

For a fraction of 6 out of 11 ART-treated PWH, experiments were performed in parallel in the presence of ARVs (integrase inhibitor Raltegravir; protease inhibitor Saquinavir), to block the infection of new cells by progeny virions produced upon TCR-mediated VR reactivation (Figure 1). Results in Figure 1B reveal, a strong decrease in integrated HIV-DNA levels mediated by ARVs in all three conditions, with the abrogation of differences observed between medium and metformin *versus* INK128 in the absence of ARVs. Similarly, in the presence of ARVs, cell-associated and soluble HIV-p24 levels became low/undetectable, with no further effects of metformin and INK128 observed under these conditions (Figure 1C-E). Thus, the proviral effects of metformin are abrogated in the presence of ARVs, which block cell-to-cell transmission of virions newly produced by reactivated VR *in vitro*. Together these results support a model in which metformin exerts its proviral effects by facilitating cell-to-cell transmission independently of cell-free virion release, consistent with previous reports^54,55^.

To get more mechanistic insights into the metformin mechanism of action, cells harvested at day 12 post-TCR triggering were analyzed for the expression of RORC2, the Th17 transcription master regulator^56^, CCR6, a Th17 cell-surface marker^57^, and IL-17A, the hallmark Th17 lineage cytokine^58^. Metformin *versus* medium increased the frequency of RORC2^+^ and CCR6^+^ T-cells and the intensity of RORC2 and CCR6 expression at single-cell level, (Supplemental Figure 2A-C).

Also, metformin *versus* medium slightly increased the frequency of CD4^+^ T-cells co-expressing RORC2 and CCR6, identified as Th17-like cells (Supplemental Figure 2A and D), but had no effect IL-17A production in cell-culture supernatants (Supplemental Figure 2E). For INK128 *versus* medium, despite an increase in the frequency of cells expressing RORC2 or CCR6 (Supplemental Figure 2A-C), there were no changes in the frequency of RORC2^+^CCR6^+^ T-cells (Supplemental Figure 2D), but a significant reduction in IL-17A production in cell-culture supernatants (Supplemental Figure 2E). Thus, metformin increased the frequency of CD4^+^ T-cells with a Th17 phenotype, without proportionally increasing their effector functions (*i.e.,* IL-17A production).

These results reveal that, in contrast to INK128 that inhibits both HIV-1 outgrowth and Th17 effector functions, metformin increases the frequency of T-cells expressing cell-associated HIV-p24 *via* mechanisms independent on free-virion release and maintains Th17 functions. These observations raise new questions on the effects of metformin on specific post-integration steps of the HIV-1 replication cycle.

### Metformin does not affect HIV-1 transcription in CD4^+^ T-cells of ART-treated PWH

Considering the capacity of metformin to increase cell-associated HIV-p24 expression in VOA (Figure 1), we further investigated the role of metformin on HIV-1 transcription. Memory CD4^+^T-cells of ART-treated PWH were stimulated *via* CD3/CD28 Abs and cultured in the presence or the absence of metformin or INK128 for 3 days. To prevent HIV-1 replication in culture, experiments were performed in the presence of ARVs (Raltegravir, Saquinavir) (Figure 2A). As expected, metformin treatment did not affect the levels of integrated HIV-DNA in the presence of ARVs (Figure 2B). Levels of cell-associated HIV-RNA, as well as the HIV-RNA/DNA ratios (surrogate maker of HIV transcription^46,59,60^) were slightly decreased in specific donors by metformin compared to the control condition, but differences did not reach statistical significance in all donors tested (Figure 2C-D). Similarly, INK128 had no effect on HIV-DNA nor HIV-RNA levels (Figure 2 B-D). These results demonstrate that, under these specific experimental settings, metformin does not exert an effect on TCR-mediated HIV-1 transcription.

**Figure 2:**
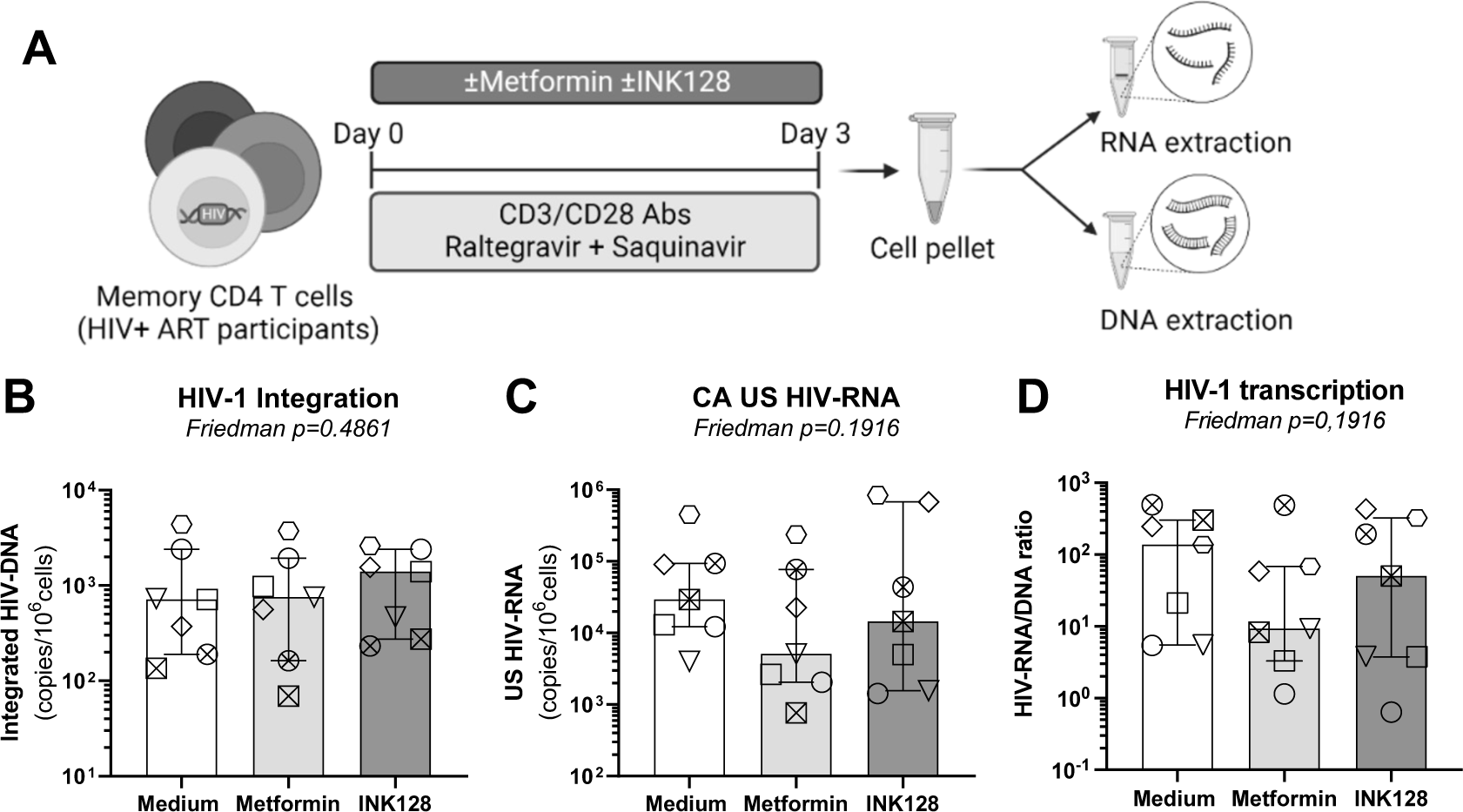
Effects of metformin on HIV-1 transcription in memory CD4^+^ T-cells of ART-treated PWH. **(A)** Shown is the experimental flow chart. Briefly, memory CD4^+^ T-cells from ART-treated PWH were stimulated by CD3/CD28 Abs in presence of ARVs (Saquinavir 5 µM, Raltegravir 200 nM) and in the presence or the absence of metformin (1 mM) or INK128 (50 nM) for 3 days. Cells were collected for dual extraction of cell-associated (CA) RNA and DNA. **(B)** Levels of integrated HIV-DNA Alu/LTR primers were quantified by nested real-time PCR and normalized per number of CD3 copies. **(C)** Levels of CA unspliced (US) HIV-RNA (Gag primers) were quantified by nested real-time RT-PCR and normalized to the number of HIV-DNA copies *per* 10^6^ cells. **(D)** The HIV RNA/DNA ratio was used as a surrogate marker of HIV-1 transcription. Each symbol represents one donor (n=7; median ± interquartile range). Friedman p-values are indicated on the graphs. Uncorrected Dunn’s multiple comparison p-values did not reach statistical significance and are not shown.

### Metformin boosts cell-associated HIV-p24 expression upon infection *in vitro*

To get insights into the molecular mechanisms underlying differences between metformin and INK128, cells were analyzed in parallel for the expression of the HIV-1 entry receptor CD4, and co-receptors CCR5 and CXCR4. Metformin did not impact on CD4 and CXCR4 surface protein expression, while INK128 slightly decreased CD4 and increased CXCR4, mainly in terms of Geometric MFI expression at single-cell level (Supplemental Figure 3A-B). Levels of CCR5 mRNA expression tended to decrease, with differences reaching statistical significance in matched comparisons for INK128 but not metformin *versus* medium (Supplemental Figure 3C).

To further document metformin effects on HIV-1 replication, TCR-stimulated CD4^+^ T-cells from HIV-uninfected participants were exposed to a replication-competent CCR5-tropic HIV_NL4.3BAL_ virus (Figure 3A). Although metformin and INK128 treatment did not impact on levels of HIV-DNA integration at day 3 post-infection (Figure 3B), both reduced HIV-p24 levels in cell-culture supernatant (Figure 3C-D). In contrast to INK128, metformin increased cell-associated HIV-p24 levels in terms of frequency of CD4^low^HIV-p24^+^ and intensity of HIV-p24 expression at single-cell level (Figure 3E-G). Metformin also significantly increased the frequency of CD4^high^HIV-p24^+^ T-cells, as well as their HIV-p24 expression at single-cell level (Figure 3E and H<u>-I</u>). These results demonstrate that metformin facilitates cell-associated HIV-p24 expression, resulting in the expansion of HIV-p24^+^ cells with a productively infected phenotype (CD4^low^) and bystander cells coated with newly formed virions (CD4^high^), likely by promoting cell-to-cell transmission of infection, *via* mechanisms independent of free-virion release.

**Figure 3:**
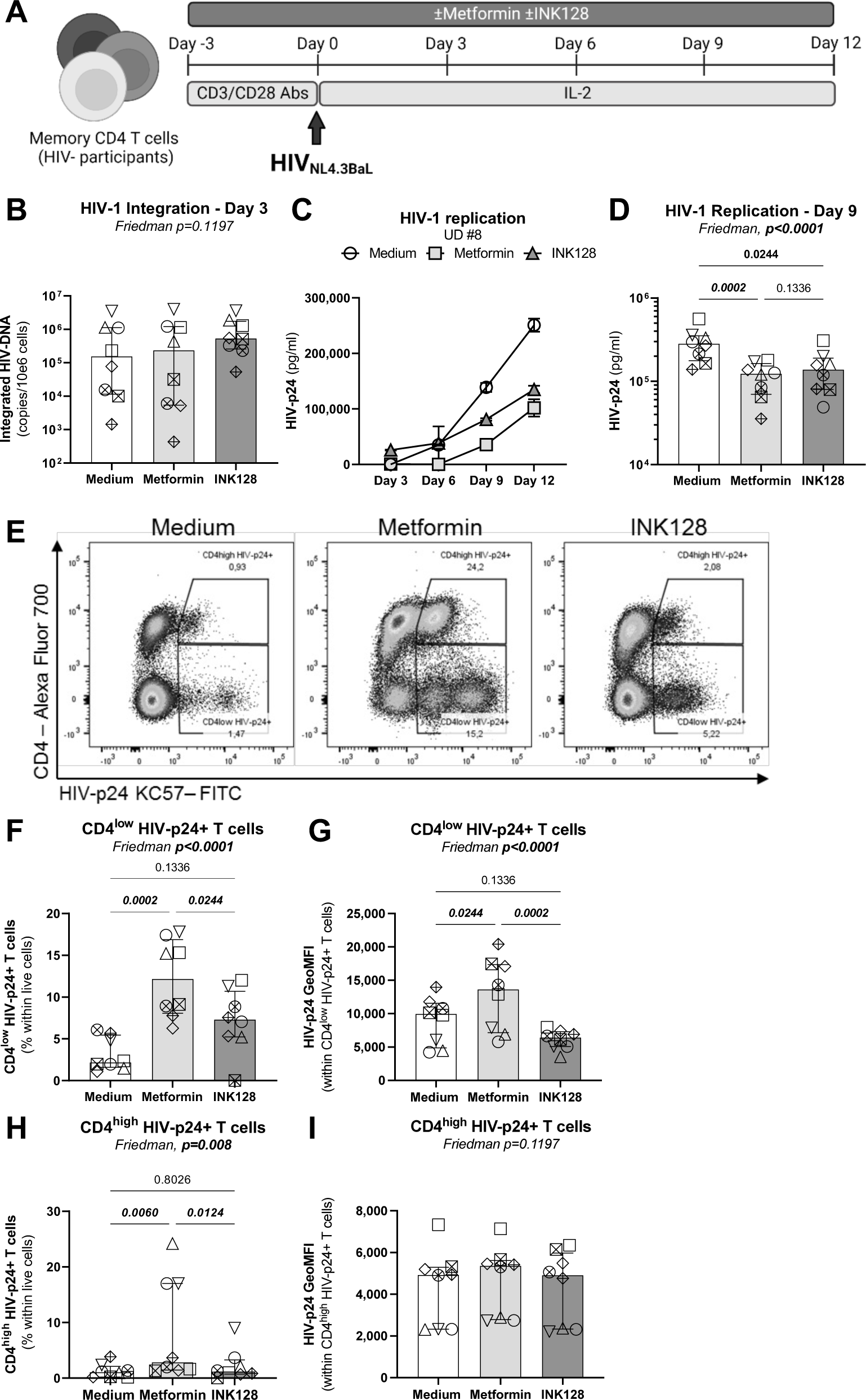
Effects of metformin on HIV-1 replication *in vitro* in memory CD4^+^ T-cells. **(A)** Shown is the flow chart for the HIV-1 infection *in vitro*. Briefly, memory CD4^+^ T-cells from HIV-uninfected donors were stimulated with anti-CD3/CD28 Abs in the absence/presence of metformin (1 mM) or INK128 (50 nM) for 3 days. Then, cells were exposed to the replication-competent NL4.3BaL HIV strain (50 ng HIV-p24/10^6^ cells). Cell-culture supernatants were collected and media containing IL-2 and/or metformin or INK128 was refreshed every 3 days until day 12 post-infection. **(B)** Integrated HIV-DNA levels were quantified by real-time nested PCR at day 3 post infection. Shown are **(C)** levels of HIV-p24 in cell-culture supernatants quantified by ELISA in one representative donor, and **(D)** statistical analysis of HIV-1 replication at day 9 post-infection in cells from n=8 different donors. Shown is **(E)** the dot plot analysis of the intracellular HIV-p24 and surface CD4 expression allowing the identification of CD4^low^HIV-p24^+^ cells (productively infected) and CD4^high^HIV-p24^+^ cells (recently infected); as well as **(F)** the statistical analysis of CD4^low^HIV-p24^+^ T-cells frequency and **(G)** the geometric MFI of HIV-p24 expression. Shown are the statistical analysis **(H)** of CD4^high^HIV-p24^+^ T-cells frequency and **(I)** the geometric MFI of HIV-p24 expression. Each symbol represents 1 donor (n=8; median ± interquartile range). Friedman and uncorrected Dunn’s multiple comparison p-values are indicated on the graphs.

Similar to results obtained in Supplemental Figure 2, metformin acted on memory CD4^+^ T-cells from uninfected participants to increase the frequency and intensity of RORC2 and CCR6 expression at single-cell level, enhanced the frequency of RORC2^+^CCR6^+^ T-cells (Supplemental Figure 4A-F), and maintained T-cell capacity to produce IL-17A in response to TCR triggering upon culture *in vitro* (Supplemental Figure 4G-H). In contrast, INK128 did not increase RORC2 expression and the frequency of RORC2^+^CCR6^+^ T-cells (Supplemental Figure 4A-F), and inhibited IL-17A production early upon TCR triggering (Supplemental Figure 4G-H). Thus, the effects of metformin on cell-associated HIV-p24 expression coincide with the promotion of a Th17 phenotype and the preservation of Th17 effector functions in memory CD4^+^ T-cells.

### Metformin facilitates HIV-1 replication post-integration and prior to viral release

To localize the step(s) of the HIV-1 replication cycle affected by metformin, single-round infection with a VSVG-pseudotyped HIV-1 was performed on memory CD4^+^ T-cells from HIV-uninfected participants in the presence/absence of metformin or INK128 (Figure 4A). In agreement with results in Figures 2-3, metformin did not significantly decrease the levels of early and late reverse transcripts, nor HIV-DNA integration (Figure 4B-D), further supporting the idea that metformin acts at post-integration level(s). In contrast, INK128 significantly reduced levels of RU5 and Gag HIV-DNA (Figure 4B-C) and tended to reduce HIV-DNA integration (Figure 4D). In this model of single-round infection, where cell-to-cell transmission does not occur given the absence of Env, metformin slightly increased the frequency of CD4^low^HIV-p24^+^ T-cells without enhancing the intensity of HIV-p24 expression *per* cell (Figure 4E-G). Of note, the CD4^high^HIV-p24^+^ T-cell population was not observed in this single-round infection system (Figure 4E), likely since Env-deficient virions cannot bind to new cells. Similar to results in Figures 2-3, metformin did not increase_the HIV-p24 release in cell-culture supernatants (Figure 4H). In contrast, INK128 diminished the intensity of HIV-p24 expression at single-cell level, although it exerted no effect on HIV-p24 levels in cell-culture supernatants (Figure 4F-H). These results suggest that metformin acts at post-integration levels by facilitating cell-associated HIV-p24 expression, with no effect on viral release.

**Figure 4:**
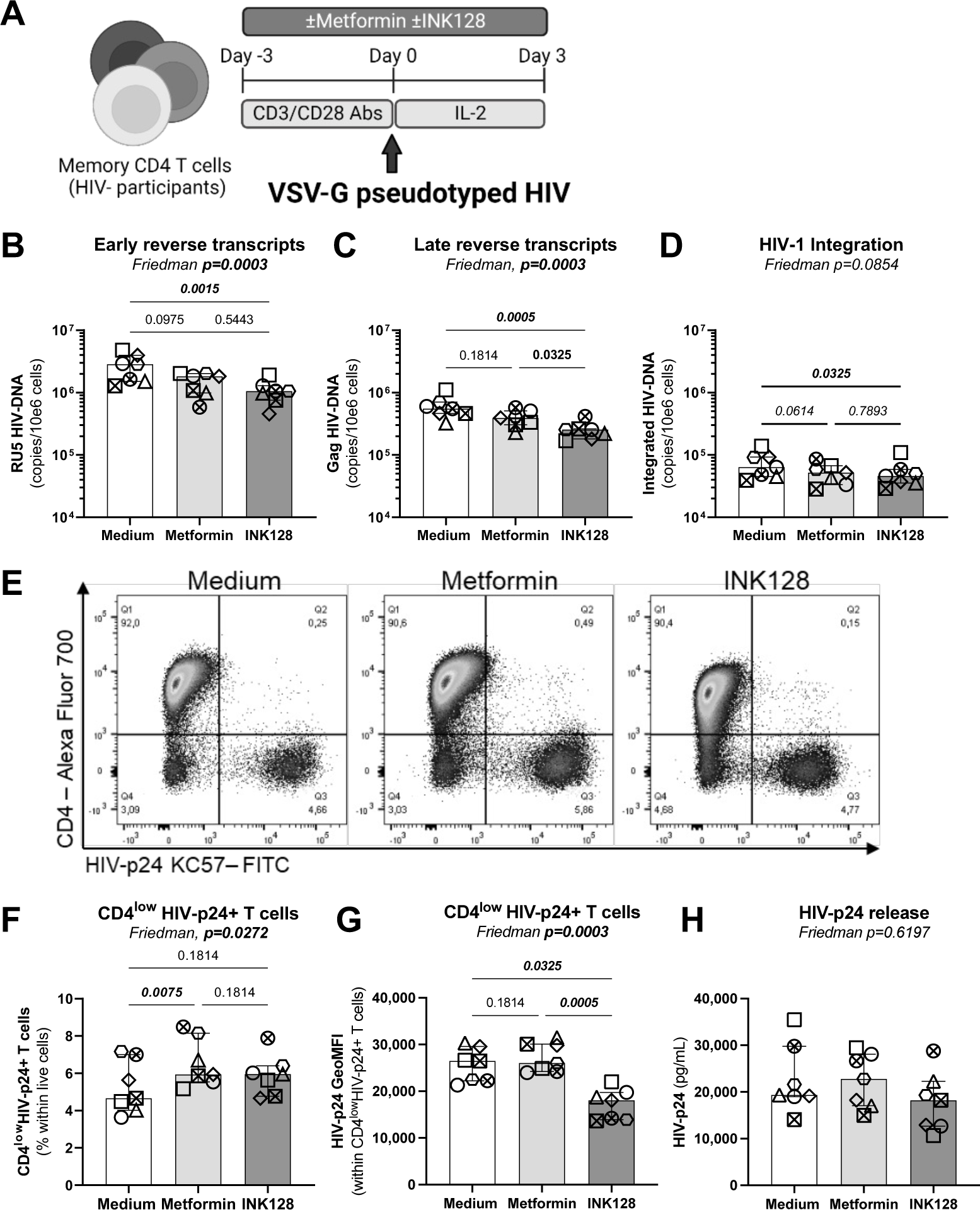
Effects on metformin on single-round HIV-1 infection *in vitro*. **(A)** Shown is the flow chart for the single-round HIV-1 infection *in vitro*. Briefly, memory CD4^+^ T-cells from HIV-uninfected donors were stimulated by anti-CD3/CD28 Abs in the absence/presence of metformin (1 mM) or INK128 (50 nM) for 3 days. Then, cells were exposed to a replication-incompetent single-round VSV-G-pseudotyped HIV-1 construct (100 ng HIV-p24/10^6^ cells). Cell-culture supernatants and cells were collected at day 3 post-infection. Shown are levels of early (RU5) **(B)** and late HIV reverse transcripts (Gag) **(C)**, as well as integrated HIV-DNA **(D)** quantified by real-time nested PCR. Shown are representative flow cytometry dot plots of intracellular HIV-p24 and surface CD4 expression from one donor **(E)** and statistical analysis of the productively infected CD4^low^HIV-p24^+^ T-cells in terms of frequencies **(F)**and the geometric MFI of HIV-p24 expression (**G)**. Shown are absolute HIV-p24 levels in cell culture supernatants quantified by ELISA (**H)**. Each symbol represents one donor (n=7; median ± interquartile range). Friedman and uncorrected Dunn’s multiple comparison p-values are indicated on the graphs.

### Metformin increases surface expression of BST2 on productively infected T-cells *in vitro*

HIV-1 release is controlled by complex mechanisms, including BST2, a protein that sequester newly formed viral particles at the cell-surface membrane^61,62^. The HIV-1 accessory protein Vpu counteracts the effects of BST2 *via* mechanisms involving BST2 downregulation^62,63^ or displacement from the site of viral assembly^64^, with BST2 mediating cell-to-cell transmission of Vpu-defective HIV-1 virions^54^. Considering the discrepancy between the effects of metformin on the frequency of productively infected cells and virion release, we hypothesized that metformin limits virion release and facilitates their cell-to-cell transmission by promoting BST2 expression. To test the possibility, memory CD4^+^ T-cells harvested at day 12 post-infection with HIV-1 _NL4.3Bal_ *in vitro* (Figure 3A) and memory CD4^+^ T-cells of ART-treated PWH harvested at day 12 post-TCR triggering (Figure 1A) were analyzed for surface expression of BST2 and cell-associated HIV-p24. Results in representative donors depicted in Figure 5A reveal the typical down regulation of BST2 on productively infected T-cells. Further, the expression of BST2 was analyzed on productively infected (CD4^low^HIV-p24^+^) *versus* uninfected (CD4^+^HIV-p24^-^) T-cells upon HIV_NL4.3BaL_ infection *in vitro* (Figure 5B-C), in VOA (Figure 5D-E), and in CD4^+^ T-cells unexposed to HIV-1 *in vitro* (Figure 5F). Upon HIV_NL4.3BaL_ infection *in vitro*, metformin and INK128 significantly increased BST2 expression at the surface of productively infected but not uninfected T-cells (Figure 5B-C). In contrast, INK128 but not metformin increased surface BST2 expression on productively infected T-cells in VOA (Figure 5D-E). The upregulation of BST2 was not observed when uninfected memory CD4^+^ T-cells were exposed to metformin or INK128 (Figure 5D).

**Figure 5:**
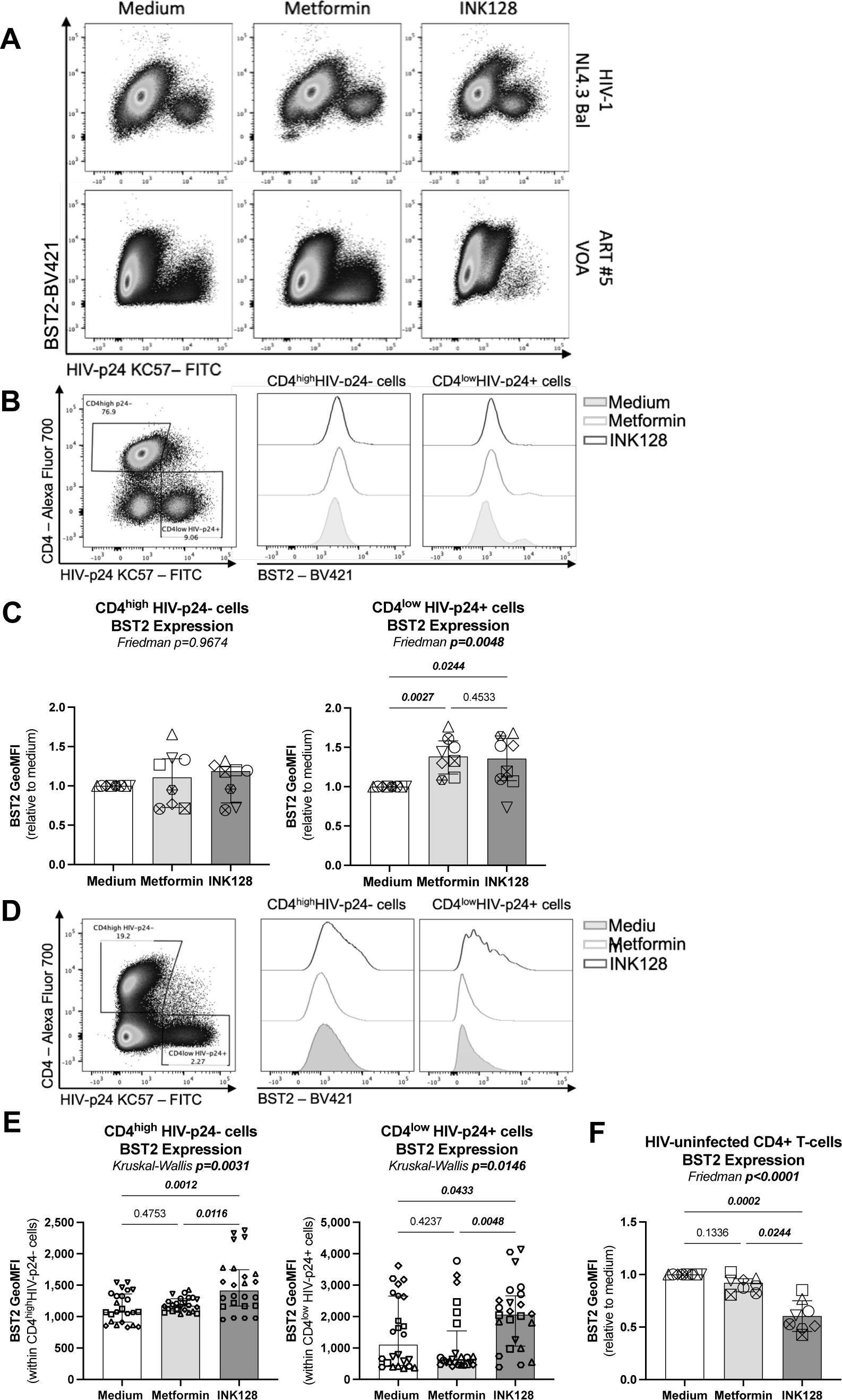
Metformin and INK128 increases BST2 expression on productively infected CD4^low^HIV-p24^+^ T-cells. **(A)** Shown are representative flow cytometry dot plots of BST2 and HIV-p24 co-expression on T-cells at day12 post-infection with HIV-1_NL4.3 Bal_ *in vitro* (upper panel) and at day 12 post TCR-mediated viral reservoir reactivation in VOA (bottom panel). **(B-C)** The HIV-1_NL4.3Bal_ infection *in vitro* was performed as described in Figure 3 <u>on cells from n=8 HIV-uninfected participants</u>. Cells collected at day 12 post-infection were stained on the surface with CD4 and BST2 antibodies and intracellularly with HIV-p24 antibodies and analyzed by flow cytometry for n=8. Shown are **(B)** levels of BST2 expression on uninfected (CD4^high^HIV-p24^-^) and productively infected (CD4^low^HIV-p24^+^) T-cells in one representative donor and **(C)** statistical analysis of BST2 expression (GeoMFI) relative to the medium condition (considered 1). **(D-E)** The VOA was performed as described in Figure 1 with cells from n=6 ART-treated PWH. Cells collected at day 12 post-stimulation were stained on the surface with CD4 and BST2 Abs and intracellularly with HIV-p24 Abs and analyzed by flow cytometry. **(D-E)** Shown are levels of BST2 expression on CD4^high^HIV-p24^-^ and CD4^low^HIV-p24^+^ T-cells in **(D)** one representative donor and **(E)** statistical analysis of BST2 expression (absolute GeoMFI). (**F)** Shown is the BST2 expression relative to the medium condition of HIV-uninfected memory CD4^+^ T-cells at day 3 post-TCR stimulation. Each symbol represents one donor (median ± interquartile range). Friedman **(C and F)**, Kuskal-Wallis **(E)** and uncorrected Dunn’s multiple comparison p-values are indicated on the graphs

These results reveal that metformin prevents the HIV-mediated downregulation of surface BST2 expression on productively infected cells, only upon exposure to HIV-1 *in vitro*, while INK128 demonstrated to be a robust modulator of surface BST2 expression in VOA as well. The fact that metformin and INK128 failed to modulate BST2 expression in the absence of HIV-1 exposure points to an HIV-dependent mechanisms of action for these two drugs.

### Metformin increases intracellular Bcl-2 expression

We previously reported that 12 weeks of metformin treatment in complement of ART in PWH increased Bcl-2, a survival marker, in colon CCR6^+^ CD4^+^ T-cells^46^. We therefore tested the possibility that metformin increases the frequency of productively HIV-infected T-cells by promoting their survival in a Bcl-2-dependent manner. Memory CD4^+^ T-cells harvested at day 12 post-infection with HIV-1 _NL4.3Bal_ *in vitro* were analyzed for intracellular Bcl-2 expression by flow cytometry on productively infected (CD4^low^HIV-p24^+^) *versus* uninfected (CD4^high^HIV-p24^-^) T-cells (Figure 6A). Metformin but not INK128 significantly increased the expression of Bcl-2 in both productively HIV-infected and uninfected CD4^+^ T-cells (Figure 6B). These results suggest that metformin treatment promotes cell survival.

**Figure 6:**
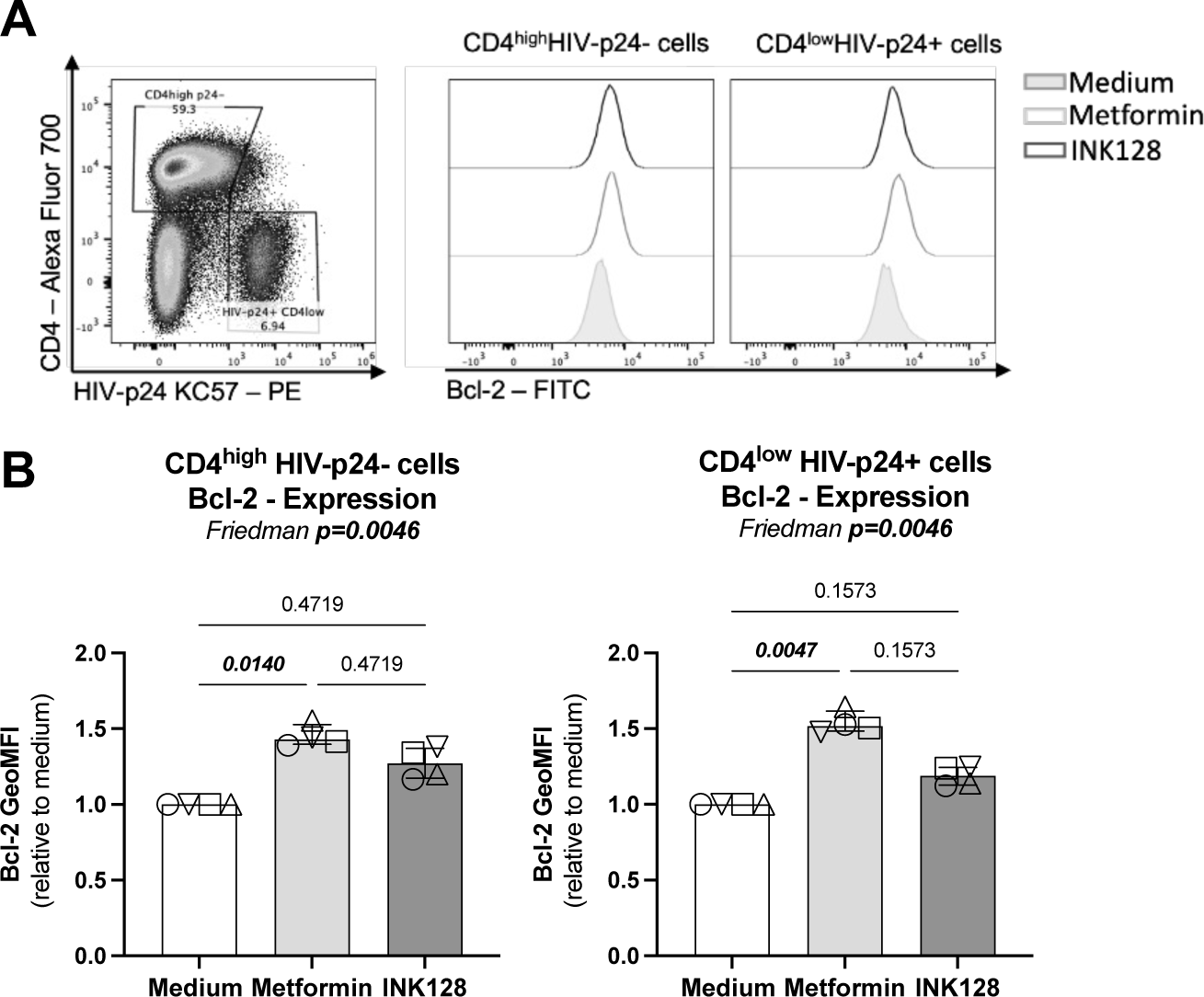
Metformin increases Bcl-2 expression on HIV-infected and uninfected T-cells. The HIV-1_NL4.3Bal_ infection *in vitro* was performed as described in Figure 3 on cells from HIV-uninfected participants. Cells collected at day 12 post-infection were stained on the surface with CD4 antibodies and intracellularly with HIV-p24 and Bcl-2 antibodies and analyzed by flow cytometry. Shown are **(A)** levels of Bcl-2 expression on uninfected (CD4^high^HIV-p24^-^) and productively infected (CD4^low^HIV-p24^+^) T-cells in one representative donor and **(B)** statistical analysis of Bcl-2 expression (GeoMFI) relative to the medium condition (considered 1). Each symbol represents one donor (n=4; median ± interquartile range). Friedman and uncorrected Dunn’s multiple comparison p-values are indicated on the graphs.

### Metformin facilitates recognition of reactivated VR by HIV envelope antibodies

Since metformin increased the frequency of CD4^low^HIV-p24^+^ T-cells upon TCR triggering of CD4^+^ T-cells from ART-treated PWH (Figure 1), we hypothesized that metformin also facilitates the recognition of reactivated VR by HIV-1 envelope (Env) Abs, thus facilitating their purging *via* Abs-dependent mechanisms *in vivo*. To test this hypothesis, we performed a VOA, as described in Figure 1A. Cells harvested at day 12 post-TCR triggering in the presence or the absence of metformin were stained on the surface with CD4 Abs and a set of broadly neutralizing (bNAbs; 2G12, PGT121, PGT126, PGT151, 3BNC117, 101074, VRC03) and non-neutralizing (nnAbs; F240, 17b, A32) HIV-1 Env Abs, and intracellularly with HIV-p24 KC57 Abs (Figure 7A).

**Figure 7:**
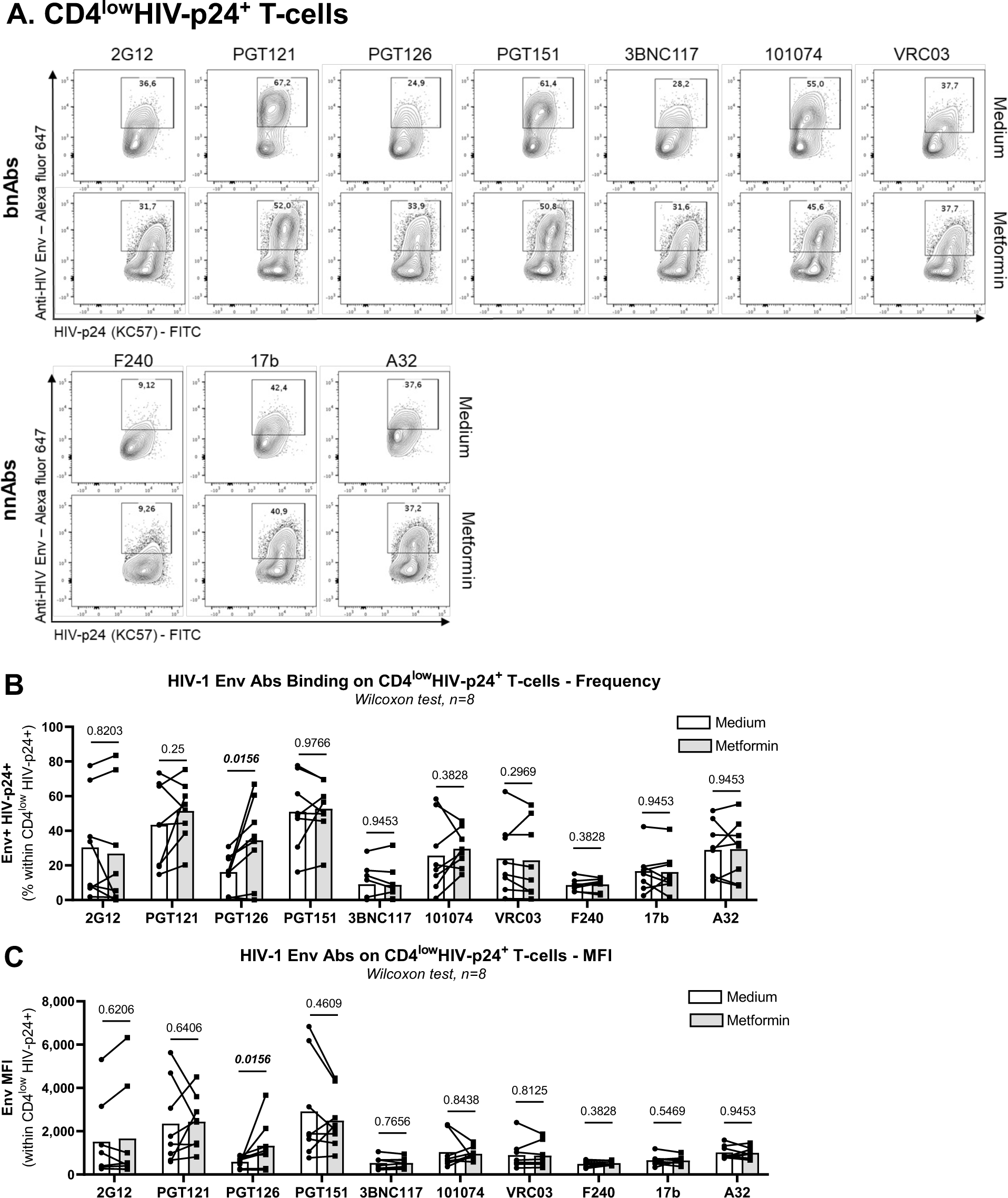
Metformin facilitates the recognition of reactivated HIV reservoirs by anti-HIV Env antibodies. The VOA was performed on memory CD4+ T-cells from ART-treated PWH, as described in Figure 1A. Cells harvested at day 12 post-TCR triggering and cultured in the presence/absence of metformin were stained on the surface with a set of unconjugated human anti-HIV-1 Env bNAbs (2G12, PGT121, PGT126, PGT151, 3BNC117, 101074, VRC03) and nnAbs (F240, 17b, A32), followed by incubation with anti-human Alexa Fluor 647-conjugated secondary Abs. Further, cells were stained on the surface with CD4 Abs, as well as intracellularly with HIV-p24 Abs. **(A)** Show are dot plot representations of HIV-p24 and HIV-Env co-expression in one representative donor. **(B-C)** Shown are statistical analysis of the frequency of anti-HIV-Env Abs binding on CD4^low^HIV-p24^+^ T-cells **(B)**, as well as the geometric MFI of anti-HIV-Env Abs binding on CD4^low^HIV-p24^+^ T-cells **(C)**. Wilcoxon test p-values are indicated on the graphs. Each symbol represents one donor (n=8; median ± interquartile range).

Different HIV-1 Env Abs showed a distinct ability to bind on productively infected cells (CD4^low^HIV-p24^+^) exposed or not to metformin (Supplemental Figure 5A). Staining performed in parallel on cells from one HIV-uninfected donor showed low/undetectable levels of non-specific Abs binding (Supplemental Figure 5B). Similar to results in Figure 1, metformin increased the frequency of T-cells recognized by both HIV-p24 Abs and specific HIV-1 Env Abs (*i.e.,* PGT121, PGT126, PGT151, 101074, F240, A32) (Supplemental Figure 5C). When the gating was performed on productively infected T-cells (CD4^low^HIV-p24^+^) (Supplemental Figure 5A), metformin increased the recognition of these cells by the PGT126 bnAbs, in terms of frequency and mean fluorescence intensity (Figure 7A-C). PTG126 bnAbs recognizes the “*closed*” conformation of the HIV-1 Env (high-mannose patch on HIV gp120)^65^, indicative that, upon TCR-mediated reactivation of VR in ART-treated PWH, metformin increased the surface expression of HIV-1 Env in its “*closed*” conformation. Whether this improved recognition could translate in potential killing of infected cells by ADCC remains to be determined.

## DISCUSSION

Although ART has saved and substantially improved the life of PWH, the treatment is not curative and chronic HIV-1 infection is associated with several comorbidities that represent a global health burden^1,11^. New therapeutic strategies are needed to reduce chronic inflammation and improve immune functions. HIV-1 infection modifies the metabolism of immune cells and thus, cellular metabolism could be a potential target for HIV-1 cure interventions^18^. In this manuscript, by using metformin, an FDA approved anti-diabetic drug that reduces mTOR pathway activity^44,66^, we expected to reduce HIV-1 replication. In contrast to our prediction, metformin treatment increased the frequency of productively infected T-cells in the context of a VOA we performed with memory CD4^+^ T-cells from ART-treated PWH, as well as upon HIV-1 infection *in vitro*. Nevertheless, metformin failed to enhance proportionally HIV-1 reverse transcription, integration, and transcription, and did not increase progeny virion release in cell-culture supernatants. These effects were associated with increased expression of BST2 and Bcl-2 on productively infected T-cells. Finally, metformin facilitated the recognition infected cells by HIV-1 Env Abs. The finding that metformin promotes the immune recognition of T-cells carrying translation-competent viral reservoirs emphasize the potential beneficial effect of metformin in accelerating the decay of viral reservoirs in ART-treated PWH in the context of efficient HIV-1 Env Abs-mediated antiviral responses.

In this study, we used three viral models that allowed us to dissect specific steps of the viral replication cycle affected by metformin. In the VOA, the outgrowth of replication-competent VR was quantified in memory CD4^+^ T-cells of ART-treated PWH upon culture *in vitro,* with the genetic features of integrated proviruses (mutations, deletions) remaining unknown. For *in vitro* infections, we used the HIV_NL4.3BaL_ molecular clone, a wild-type replication-competent CCR5-tropic HIV-1, and the single-round HIV_VSV-G_ construct contained a EGFP gene inserted in the Env gene, leading to the generation of Env-defective HIV-1 virions. In all these three models, metformin treatment did not increase viral release, but increased the frequency of productively infected CD4^low^HIV-p24^+^ T-cells. Metformin effects were less pronounced in experiments performed with HIV_VSV-G_, in line with the inability of Env-defective viruses to infect new cells. Metformin effects in VOA were also abrogated by ARVs, supporting the possibility of a metformin-mediated mechanism of cell-to-cell transmission independent of free virion release. These findings suggest that metformin facilitates the expression of HIV-p24 protein in reactivated viral reservoir cells.

Metformin is an indirect regulator of mTOR, which controls Th17 polarization and functions^67,68^. Direct mTOR inhibitors such as INK128 and rapamycin reduce IL-17A production^69,70^. In the literature, metformin was reported to reduce IL-17A production and RORC2 expression *in vitro*^71^. The latter report contrasts with results from our study, likely because the latter study was performed under Th17-polarizing conditions. Our results show that metformin treatment, in contrast to INK128, promotes CCR6 and RORC2 protein expression and maintains the Th17 cell effector functions (*i.e.,* IL-17A). Th17 cells are largely depleted after HIV-1 infection and their maintenance is linked to a better control of HIV-1 replication in elite controller.^72,70^ Furthermore, studies by our group demonstrated that Th17 cells are highly susceptible to HIV-1 infection given their unique high metabolic activity and transcriptional profiles.^20,22,73–76^ The pleiotropic effects of metformin on various steps of the viral replication cycle coincided with an increased Th17-polarisation phenotype (CCR6^+^RORC2^+^), and a preserved IL-17A production. Therefore, metformin treatment could exert its proviral activities by boosting Th17 polarization.

Our results on metformin-mediated decrease in viral release, with a concomitant increase in cell-associated HIV-p24, prompted us to investigate the effects of metformin on the expression of BST2/Tetherin, a host-cell restriction factor originally reported to tether progeny virions on the cell surface, thus preventing their release^62^. Further studies reported that basal levels of BST2-mediated virion tethering are required for efficient cell-to-cell transmission of HIV-1^55^, mainly in primary cells^77^. Indeed, we found that, upon HIV-1 exposure *in vitro,* metformin reduced cell-free virion levels, while enhancing the frequency of productively infected cells and boosting their BST2 expression. This evidence supports a model in which metformin limits virion release but facilitates cell-to-cell transmission by modulating the surface expression of BST2. The antiviral features of BST2 are regulated *via* glycosylation and intracellular trafficking^78,79^. Moreover, BST2 exists in two isoforms, long (L-tetherin) and short (S-tetherin), with Vpu mainly targeting the long isoforms^80^. Furthermore, the fact that BST2 acts through an interaction with the HIV-1 Env^81,62^, explains the accumulation of cell-associated HIV-p24 in T-cells upon exposure to wild type HIV_NL4.3 Bal_ but not Env-deficient VSVG-pseudotyped HIV-1. The HIV-1 accessory protein Vpu facilitates viral release by decreasing BST2 expression and its restriction activity^62^, pointing to the possibility that metformin counteracts the Vpu-mediated BST2 downregulation in infected T-cells. The ability of Vpu to counteract BST2 depends on its serine phosphorylation ^82^, a process likely modulated by metformin in mTOR-dependent manner. In VOA, BST2 expression on productively infected cells was not influenced by metformin, consistent with the fact that a metformin-mediated increase in cell-associated HIV-p24 expression was not observed at single cell level in the VOA.

This raises new questions on the specificity of metformin action in relationship with the particularities of reactivated proviruses in PWH. Molecular mechanisms by which metformin regulates BST2 expression and functions (*e.g.,* transcription of specific isoforms, glycosylation, cellular localization, Vpu-interactions) remain to be further elucidated.

The effects of metformin on the expansion of productively infected T-cells upon infection *in vitro* were associated with an increased expression of Bcl-2, a mitochondrial protein associated with cell survival. The upregulation of Bcl-2 by metformin was also observed on colon-infiltrating CCR6^+^CD4^+^ T-cells in our pilot clinical trial performed on ART-treated PWH^46^. This is consistent with the knowledge that metformin is used as an anti-aging medicine^83^. Most recent studies demonstrated the clinical benefits of Bcl-2 inhibitors (*i.e.,* Venetoclax) in promoting VR purging^84–87^. Whether metformin supplementation of ART may render reactivated VR more sensitive to Bcl-2 blockade, requires investigations in animal models and human clinical trials.

In HIV eradication strategies, both the “*shock*” and “*kill*” arms will be required^88,89^. BST2 acts as an innate sensor of viral assembly^90^, suggesting that metformin may facilitate VR sensing by the immune system *via* BST2-dependent mechanisms. Indeed, multiple studies by our group and others documented the ability of BST2 to facilitate ADCC^91–96^. In this context, we assessed the impact of metformin on the recognition of reactivated viral reservoirs by bNAbs and nnAbs. We demonstrated that metformin treatment increased the frequency of productively infected CD4^+^ T-cells recognized by the bNAbs PGT126, as well as its binding intensity at the single-cell level. The antiviral activities of PGT126 bNAbs were tested in a rhesus macaque infection model, in which PGT126 Abs administered before vaginal or rectal SHIV challenge displayed protective effects against infection acquisition^97,92^. Whether metformin can increase the ADCC activity of PGT126 Abs remains to be demonstrated. If so, a combination therapy including metformin and bNAbs could be beneficial to reduce the size of HIV-1 reservoirs during ART.

In conclusion, our results support a model in which metformin supplementation of ART acts on T-cells carrying VR to boost the expression of HIV-p24, tether the progeny virions on the cell surface, and promote their recognition by HIV-1 Env nNAbs. Considering its pleiotropic pro/antiviral effects on specific steps of the HIV-1 replication cycle, long-term double blind clinical trials should be performed to test metformin together with HIV-1 Env bNAbs in complement of ART in PWH as a novel HIV-1 remission/cure strategy to target HIV-1 reservoirs.

## LIMITATIONS

First, the metformin concentration used in our *in vitro* study was 1 mM a concentration that may not reflect the actual concentration in tissues upon metformin administration in clinic^46^. Since metformin, taken orally, mostly acts in tissues such as the liver and intestines, we cannot be sure that the effects observed *in vitro* on peripheral blood CD4^+^ T-cells reflect the reality *in vivo*. However, our decision to use metformin at 1mM is justified by its effect on mTOR activation and HIV-1 replication, without affecting cell viability and proliferation (Supplemental Figure 1). Also, studies by other groups also reported results using the same concentration of metformin^25,71^. Since metformin did not boost HIV transcription in our *ex vivo* experimental settings, further investigations are needed to decipher the molecular mechanisms used by metformin to reactivate HIV-1 from latency, likely by acting at translational level.

Moreover, the flow cytometry staining of cell-associated HIV-p24 did not allow to distinguish whether virions were trapped in the cytoplasm, at the inner or at the outer cell surface membrane. Further investigation using microscopy visualization should be performed.

Furthermore, in addition to ADCC mediated by NK cells, CD8^+^ T-cells are also key effectors for the control HIV-1 replication^98,99^. Of interest, CD8^+^ T-cells differentiation and antiviral functions are dependent on the mTOR activity^100^. In this context, the role of metformin treatment on CD8^+^ T-cell-mediated killing of HIV-infected T-cells remain to be elucidated. In line with this possibility, studies in tumor cell lines demonstrated that metformin increased the cytotoxic activities of CD8^+^ T-cells against cancerous cells^101^. Of note, eight weeks of metformin treatment in nondiabetic PWH increased the cytotoxic response of CD8^+^ T-cells^102^. These data support the idea that metformin could have a beneficial effect both on HIV-1 reservoir reactivation and on the quality of HIV-specific CD8^+^ T-cell responses.

Finally, the cohort of ART-treated PWH used in the present study was composed of a majority of Caucasian male participants (*i.e,* 1 female, 12 males; 1 Latin-American, 12 Caucasians), infected by the HIV-1 clade B. Of note, metformin treatment was not tested before on another HIV-1 clade. Differences in sex, as well as the ethnicity, could have an impact on the effect of metformin treatment considering that non-AIDS comorbidities and metabolic disorders vary depending on sex and ethnicity^103^. These aspects should be further tested in an effort to implement precision medicine strategies.

## AUTHOR CONTRIBUTIONS

Conceptualization, A.Fe, D.P. and P.A.; Methodology, A.Fe, D.P., J.R., A.Fi. and P.A.; Investigation and Formal Analysis, A.Fe, J.R. and L.R.M.; Resources, J-P.R., N.C, P.A. and A.Fi.; Writing – Original draft, A.Fe.; Writing – Review & Editing, A.Fe., D.P., J.R, A.Fi, N.C., L.R.M., J-P.R. and P.A.; Supervision A.Fi, N.C. and P.A; Funding Acquisition, P.A., Project Administration, P.A.

## Supporting information

Key ressource Table

## ACKNOWLEDGEMENTS

The authors thank Philippe St Onge and Dr. Gael Duluth (Flow Cytometry Core Facility, CHUM-Research Center, Montréal, QC, Canada) for expert technical support with flow cytometry analysis; Amelie Pagliuzza (CHUM-Research Center, Montréal, QC, Canada) for expert technical help in HIV transcription quantification; Dr. Olfa Debbeche and Laurent Knaffo (Biosafety Level 3 Core Facility CHUM-Research Cente, Montréal, QC, Canada) for assistance in manipulating infectious samples; Mario Legault (FRQ-S/AIDS and Infectious Diseases Network; Montréal, QC, Canada) for help with ethical approvals and informed consents; Josée Girouard and Angie Massicotte (McGill University Health Centre, Montréal, QC, Canada) for their key contribution to blood collection and clinical information from PLWH and uninfected study participants. The authors also thank Dr. Dana Gabuzda (Dana-Farber Cancer Institute, Boston, Massachusetts, USA), Dr Roger J Pomerantz (Thomas Jefferson University, Philadelphia, Pennsylvania, USA), and Dr. Michel Tremblay (Université Laval, Quebec, QC, Canada) for providing us with VSV-G and HIV plasmids. We thank the following collaborators for kindly providing plasmids to produce HIV-1 Env antibodies: James Robinson (Tulane University) for A32; John Mascola (Vaccine Research Center, NIAID) for VRC03; Michel Nussenzweig for 3BNC117 and 10-1074; the NIH AIDS Reagent Program for F240, 2G12 and 17b; and the International AIDS Vaccine Initiative (IAVI) for PGT121, PGT126, PGT151. Finally, the authors acknowledge the key contribution of all study participants for their precious gift of leukapheresis essential for this study.

## STAR METHODS

The source and catalogue numbers from all reagents were included in the key resource table.

### Ethics statement

Study participants were recruited at the Montreal Chest Institute, McGill University Health Centre, and Centre Hospitalier de l’Université de Montréal (Montreal, Québec, Canada), in compliance with the principles included in the Declaration of Helsinki. This study received approval from the Institutional Review Board (IRB) of the McGill University Health Centre and the CHUM Research Centre, Montreal, Quebec, Canada. All participants signed a written informed consent and agreed with the publication of the results.

### Study participants

This study was performed using Peripheral Blood Mononuclear Cells (PBMCs) from ART-treated PWH (n=13) and HIV-uninfected (n=15) study participants. PBMCs were isolated by gradient density centrifugation from leukapheresis and maintained frozen in liquid nitrogen until use, as previously described ^104^. Clinical parameters of study participants are included in Table 1 for PWH and Supplemental table 1 for HIV-uninfected donors.

**Table 1:** Clinical parameters of ART-treated PWH participants.

| ID | Sex | Ethnicity | Age<br>& | CD4<br># | CD8<br># | Plasma<br>VL* | Time since<br>infection** | ART<br>regimen | Time on<br>ART** |
| --- | --- | --- | --- | --- | --- | --- | --- | --- | --- |
| ART #1 | M | Caucasian | 36 | 542 | 803 | <40 | 13 | Stribild | 12 |
| ART #2 | M | Caucasian | 49 | 458 | 899 | <40 | 201 | Truvada<br>Viramune | 201 |
| ART #3 | M | Caucasian | 58 | 546 | 1,116 | <40 | 408 | Atripla | 369 |
| ART #4 | M | Caucasian | 44 | 546 | 775 | <40 | 154 | Complera | 25 |
| ART #5 | M | Caucasian | 51 | 546 | 1,322 | <40 | 149 | Sustiva<br>Truvada | 149 |
| ART #6 | M | Caucasian | 33 | 546 | 854 | <40 | 89 | Stribild | 77 |
| ART #7 | M | Caucasian | 21 | 546 | 399 | <40 | 8 | Stribild | 4 |
| ART #8 | M | Caucasian | 47 | 546 | 1,156 | <40 | 182 | Atripla | 63 |
| ART #9 | M | Caucasian | 59 | 546 | 836 | <40 | 273 | Symtuva | 273 |
| ART #10 | M | Caucasian | 64 | 546 | 620 | <40 | 186 | Triumeq | 186 |
| ART #11 | M | Latino-American | 31 | 546 | 1,000 | <40 | 69 | Complera | N/A |
| ART #12 | M | Caucasian | 30 | 546 | 605 | <40 | 80 | Stribild | 77 |
| ART #13 | F | Caucasian | 31 | 546 | 445 | <40 | 212 | Viracept<br>Truvada | 187 |

ID, participant identification code; ART, antiretroviral treated PWH; M, male; F, female; &, years; #, counts of cells/ $\mu$ L blood; \*, HIV-RNA copies per ml of plasma; \*\*, Months; N/A, not available
ID, participant identification code; ART, antiretroviral treated PWH; M, male; F, female; &, years;

### Memory CD4^+^ T-cell sorting

Memory CD4**^+^** T-cells were isolated from PBMCs of HIV-uninfected and ART-treated PWH by negative selection using the EasySep Human Memory CD4^+^ T Cell Enrichment Kit (StemCell Technology), following the manufacturer recommendation. The cell purity after sorting was >95%, as determined upon staining with CD3, CD4, CD45RA and CD8 Abs and flow cytometry analysis (BD LSRII).

### Flow cytometry analysis

For surface staining, cells were incubated for 30min at 4°C in PBS 1X buffer containing 10% FBS (Wisent; Cat. Num.: 091-150), 0.02% NaN3 and fluorescence-conjugated antibodies against CD3, CD4, CD8, CCR6, CD45RA, CXCR4 and BST2 (Supplemental Table 1), using a protocol we previously reported ^20,76^. Live/Dead Fixable Aqua Dead cells stain Kit was used to exclude dead cells (Invitrogen). Intracellular/nuclear staining was performed using the FoxP3 transcription factor staining buffer kit (eBioscience) and fluorescence-conjugated antibodies against HIV-p24 KC57, RORC2 and Bcl-2 (<u>Key Resources Table</u>). Flow cytometry acquisition of stained cells was performed on a BD LSRII cytometer. Flow cytometry analysis was performed using the BD Diva and FlowJo version 10. The positivity gate for RORC2 were placed using the fluorescence minus one (FMO) strategy, as reported ^105^). The positivity gate for HIV-p24 (KC57) were placed using uninfected memory CD4^+^ T-cells.

### Cell culture and activation

For TCR triggering of primary memory CD4^+^ T-cells, cells were cultured in RPMI1640 (GIBCO) cell-culture media (10% FBS, 1% Penicillin/Streptomycin (GIBCO) at 1×10^6^ cells/ml in the presence of immobilized CD3 antibodies (1 µg/ml; BD Biosciences) and soluble CD28 antibodies (1 µg/ml; BD Biosciences).

### Compounds

The following drugs were used to treat primary CD4^+^ T-cells: metformin (0.1; 0.5; 1 and 5 mM) (1,1-Dimethylbiguanide, Hydrochloride; Cat. Num. sc-202000; Santa Cruz); INK128 (50 μM) (Item No. 11811; Cayman); Saquinavir (5 μM) (NIH HIV Reagent Program), and Raltegravir (0.2 μM) (NIH HIV Reagent Program).

### HIV viral stocks

In this study, the following HIV-1 viruses were used *(i)* replication-competent CCR5 using (R5) NL4.3BAL and *(ii)* single-round VSVG-HIV-GFP, an *env*-deficient NL4.3 provirus pseudotyped with the VSV-G envelope and encoding for *gfp* in place of *env*. The NL4.3BaL HIV plasmid was provided by Michel Tremblay, Université Laval, Quebec, Canada, originating from Roger J. Pomerantz, Thomas Jefferson University, Philadelphia, PA. The plasmid pHEF Expressing Vesicular Stomatitis Virus (VSV-G) (ARP-4693) was obtained through the NIH HIV Reagent Program, Division of AIDS, NIAID, NIH, contributed by Dr. Lung-Ji Chang. The HIV vector containing the NL4-3 backbone encoding for enhanced green fluorescent protein (EGFP) in place of the Envelope (Env) (NL4.3EGFPΔEnv) was obtained through the NIH HIV Reagent Program, Division of AIDS, NIAID, NIH, contributed by Dr. Haili Zhang, Dr. Yan Zhou and Dr. Robert Siliciano. The plasmids were amplified upon bacterial transformation by MiniPrep (Promega) and MaxiPrep (Qiagen) following the manufacturer recommendation. The plasmid NL4.3Bal HIV-1 was transfected in 293T cells in order to produce the CCR5-tropic replication-competent NL4.3Bal HIV-1 viral stock. The plasmids VSV-G and NL4-3 ΔEnv EGFP were transfected together in a ratio 1:3 in 293T-cells. To perform the transfection in 293T-cells, using the X-tremeGENE 9 kit (Roche), according to manufacturer’s recommendation. Cell-culture supernatant containing the virus was collected 72h post-transfection. The NL4.3Bal HIV stock obtained on 293T-cells was passed once on TCR-activated memory CD4**^+^** T-cells and the cell-culture supernatant was collected at day 12 post-infection. The HIV viral stocks were quantified by HIV-p24 ELISA and the quantity needed for optimal infection was determined by titration on TCR-activated memory CD4**^+^** T-cells.

### HIV-1 infection in vitro

HIV-1 infection *in vitro* was performed as we previously reported ^76,106^. Memory CD4^+^ T-cells were stimulated by CD3/CD28 Abs for 3 days prior infection. Cells were exposed to NL4.3BAL (50 ng HIV-p24/10^6^ cells) for 3 hours at 37 °C and homogenized every 30 min, or VSVG-HIV-GFP (100 ng HIV-p24/10^6^ cells) and spinoculated for 1h at 300 g at room temperature. Unbound virions were removed by extensive washing with RPMI1640 10%FBS, 1%PS. Cells were cultured in the presence of IL-2 (5 ng/mL; R&D Systems) at 37°C for 12 and 3 days for NL4.3BAL and VSVG-HIV-GFP, respectively. A fraction of cells collected at day 3 post-infection was used for nested real-time PCR quantification of HIV-DNA. Cell-culture supernatants were harvested and productive infection was measured by HIV-p24 ELISA, using homemade Abs, as previously described ^107,108^, and flow cytometry analysis upon surface CD4 and intracellular HIV-p24 staining. Productively infected T-cells were identified based on their CD4^low^HIV-p24^+^ phenotype, with CD4 downregulation being indicative of productive infection, as previously reported ^109,53,110^.

### Viral Outgrowth Assay (VOA)

To measure replication-competent viral reservoirs, a simplified viral outgrowth assay was performed using a protocol developed in our lab ^51^. Succinctly, memory CD4^+^ T-cells from ART-treated PWH were cultured at 1×10^6^cells/well in RPMI1640, 10%FBS, 1% PS cell-culture media in 48-well plates in the presence of immobilized CD3 Abs and soluble CD28 Abs (1 µg/ml). At day 3, cells were washed to remove the CD3/CD28 Abs. Cells from each well were split into two new wells for optimal cell density (typically 1-2×10^6^ cells/ml/well) and cultured in the presence of IL-2 (5 ng/mL). The splitting procedure was repeated with media being refreshed every 3 days. The VOA was performed in the presence or the absence of metformin (1 mM), INK128 (50 μM) and antiretroviral drugs (ARVs; Saquinavir at 5 μM and Raltegravir at 0.2 μM). At day 12, one original replicate generated 8 splitting replicates, from which cells were harvested for the quantification of intracellular HIV-p24 expression by flow cytometry, as well as HIV-DNA by PCR. Cell-culture supernatants were collected every 3 days for HIV-p24 level quantification by ELISA.

### Quantification of early, late reverse transcript and integrated HIV-DNA

Early and late HIV-1 reverse transcripts, as well as integrated HIV-DNA levels were quantified using specific primers and probes (Supplemental Table 2), as we previously described ^20,111,112^. Briefly, cell lysates, generated by proteinase K digestion, were used to quantify HIV-DNA copies. Early reverse transcripts were amplified using primers specific for the RU5 region of the HIV genome, using SYBR green real-time nested PCR (Qiagen). Gag and integrated HIV-DNA, as well as CD3 DNA (used to normalize HIV-DNA expression per number of cells) were amplified using specific primers (Supplemental Table 2) and nested real-time PCR. The first PCR round performed with both HIV and CD3 primers was followed by a second round of PCR performed with specific internal primers and probes on the LightCycler 480II (Roche) (Supplemental Table 2). Results are expressed as HIV-DNA copies per million cells, upon normalization to CD3 copies. For all PCR quantifications, ACH2 cells (NIH HIV Reagent Program) carrying one copy of integrated HIV-DNA per cells, were used as a standard curve.

### Quantification of HIV-1 transcription

The HIV-1 RNA/DNA ratios were used as surrogate markers of HIV transcription, as we and other groups previously reported ^46,59,60^. Memory CD4^+^ T-cells from ART-treated PWH, cultured in five replicates at 1×10^6^cells/well in RPMI1640, 10% FBS, 1% PS media in 48-well plates, were stimulated via CD3/CD28 Abs in presence/absence of metformin (1 mM) or INK128 (50 μM) in addition with ARVs (Saquinavir (5 μM), and Raltegravir (0.2 μM)). At day 3 post-TCR triggering, cells were harvested, washed and replicates were pooled. Cell-associated (CA) RNA and DNA were dually extracted using the AllPrep DNA/RNA/miRNA Universal Kit (Qiagen). The quantity and the quality of extracted RNA/DNA were evaluated by Nanodrop. For HIV-RNA quantification, in a first step, CA HIV-RNA were reverse transcribed and amplified by RT-PCR using external primers annealing the LTR Gag region (Supplemental Table 3) and SuperScript III One-Step RT-PCR Taq polymerase (Invitrogen). In a second step, PCR amplification was performed with internal primers and PerfeCTa qPCR ToughMix Low ROX (QuantaBio), using the RotorGene Instrument (Supplemental Table 3). A standard curve was generated using a plasmid-based transcription *in* vitro containing LTR-Gag (pIDT-Blue) (provided by Dr Nicolas Chomont, Université de Montréal, Québec, Canada). For HIV-DNA quantification, in a first step, HIV-DNA was amplified using external primers recognizing the HIV-LTR Gag region and the Alu region. In the integrated HIV-DNA PCR, specific primers for CD3 were added to allow normalization on the number of cells within samples. In a second step, HIV-DNA and CD3 DNA were amplified separately using specific primers and probes (Supplemental Table 3) on the RotorGene PCR machine. CA DNA extracted from ACH2 cells was used as a standard curve. All measures were performed in triplicate. Results are expressed as HIV-RNA and HIV-DNA copies per 10^6^ cells.

### Anti-HIV-1 envelope antibodies recognition of productively HIV-infected T-cells

Memory CD4^+^ T-cells harvested at Day 12 of VOA were analyzed by flow cytometry for the binding of a panel human Abs directed against the HIV-1 Env. The following antibodies were used: anti-gp41 F240; anti-cluster A A32, anti-coreceptor binding site 17b; anti-CD4 binding site VRC03, 3BNC117; anti-gp120 outer domain 2G12; the gp120-gp41 interface PGT151 and anti-V3 glycan PGT121, PGT126, 101074. The goat anti-human IgGs conjugated with Alexa Fluor 647 (Invitrogen) were used as a secondary Abs to determine the levels of anti-HIV-gp120 Abs binding. Then the cells were stained on the surface with CD4-Alexa Fluor 700 Abs and intracellularly with HIV-p24-FITC Abs (<u>Key resource Table</u>). Productively infected T-cells were identified based on their CD4^low^HIV-p24^+^ phenotype. The viability dye was used to exclude dead cells from the analysis.

### HIV-1 Env Antibody production

FreeStyle 293F cells (Thermo Fisher Scientific) were grown in FreeStyle 293F medium (Thermo Fisher Scientific) to a density of 1 × 10^6^ cells/mL at 37°C with 8% CO_2_ with regular agitation (150 rpm). Cells were transfected with plasmids expressing the light and heavy chains of A32 (kindly provided by James Robinson); VRC03 (kindly provided by John Mascola); 3BNC117, 10-1074 (kindly provided by Michel Nussenzweig); F240, 2G12, 17b (NIH AIDS Reagent Program); PGT121, PGT126, PGT151 (IAVI), using ExpiFectamine 293 transfection reagent, as directed by the manufacturer (Thermo Fisher Scientific). One week later, the cells were pelleted and discarded. The supernatants were filtered (0.22-μm-pore-size filter), and antibodies were purified by protein A affinity columns, as directed by the manufacturer (Cytiva, Marlborough, MA, USA).

### Western Blot

The visualization of total and phosphorylated mTOR and S6 ribosomal proteins was performed using protocols established in the laboratory, as we previously reported ^20^. Cells were lysed with RIPA buffer (Cell Signaling) containing phosphatase inhibitors and protease inhibitors (Milipore Sigma) for 5 min at 4°C and sonicated 3 times for 5 seconds on ice. Lysed pellets were centrifuged at 14,000 g for 10 min to remove cell debris. Proteins were quantified using the kit DM^TM^ Protein Assay (Bio-Rad). Loading of proteins (10 μg/well) was performed onto a 7% acrylamide SDS gel for mTOR and 15% for S6 ribosomal and electrophoretic migration was performed (1h10 at 150V). Migrated proteins were transferred by electrophoresis on activated PVDF membrane (1h at 100V). Membranes were blocked for 45min at room temperature with TBST 5% BSA, 0.1% Tween buffer. To measure phosphorylated proteins, membranes were bloated with primary Abs against Phosphorylated Ribosomal S6 (EMD Milipore;) and Phosphorylated mTOR (Cell Signaling) Abs overnight at 4 °C. Then membranes were washed with TBST 0.1 %Tween buffer and incubated with secondary antibody anti-Rabbit IgG HRP-linked (Cell Signaling) for one hour at room temperature. Proteins were revealed with Clarity Max™ Western ECL Substrate, (Bio-Rad). For total mTOR and S6, the same membranes were stripped with Re-Blot Plus Strong Solution (EMD Milipore) and re-bloated with the appropriate primary and secondary Abs. For β-actin, the same membranes were stripped with Re-Blot Plus Strong Solution (EMD Milipore) and re-bloated with the primary anti-β-actin Abs (Sigma Aldrich) and HRP conjugated-secondary Abs (Invitrogen).

### Statistical analysis

Statistical analyses were performed with GraphPad Prism 9.0.1. Statistical tests used are indicated in the figure legends and p-values are indicated on the graphs. P-values ≤ 0.05 were considered statistically significant.

## SUPPLEMENTAL FIGURE LEGENDS

**Supplemental Figure 1: Effects of metformin on mTOR activation, viability, and proliferation in memory CD4^+^ T-cells.** Memory CD4^+^ T-isolated from PBMCs of HIV-uninfected donors were stimulated by anti-CD3/CD28 Abs for 3 days in the presence/absence of different doses of metformin (0.1, 0.5, 1 and 5 mM). **(A-D)** Cell lysates were used to measure mTOR activation by visualizing the expression of phosphorylated mTOR (A-B) and S6 ribosomal protein (C-D) by western blotting. Shown is the mTOR and S6 ribosomal protein bands (A and C), as well as the quantification of the mTOR and S6 ribosomal protein bands relative to β-actin in cells from one representative donor. **(E-F)** The cell viability and proliferation were evaluated by cytometry upon staining with the viability dye Aqua Vivid and intranuclear staining with the Ki67 Abs, respectively. Shown is cell viability **(E)** and proliferation **(F)** in experiments performed with cells from n=8 different HIV-uninfected participants. Friedman and uncorrected Dunn’s multiple comparison p-values are indicated on the graphs.

**Supplemental Figure 2: Metformin increases the expression of RORC2 and CCR6 on memory CD4^+^ T-cells from ART-treated PWH.** The VOA was performed on memory CD4^+^ T-cells from ART-treated PWH, as described in Figure 2A. Shown are representative flow cytometry dot plots of intracellular RORC2 and surface CCR6 expression **(A)**; as well as the statistical analysis of the RORC2^+^ T-cell frequency **(B, left panel)** and the geometric MFI of RORC2 expression **(B, right panel);** the frequency of CCR6^+^ T-cells **(C, left panel)** and the geometric MFI of CCR6 expression **(C, right panel)**; and the frequency of Th17-like cells identified as cells with a CCR6^+^RORC2^+^ phenotype **(D)**. (Finally, shown is the statistical analysis of IL-17A production quantified in the cell-culture supernatant by ELISA **E)**. Each symbol represents one donor (n=11 without ARVs); bars indicate the median ± interquartile range. Kruskal-Wallis test and uncorrected Dunn’s multiple comparison p-values are indicated on the graphs.

**Supplemental Figure 3: Metformin does not impact the expression of CD4, CXCR4 and CCR5.** Memory CD4^+^ T-cells from HIV-uninfected donors were stimulated by anti-CD3/CD28 Abs in the presence/absence of metformin (1 mM) or INK128 (50 nM) for 3 days. The expression of CD4 and CXCR4 was evaluated by flow cytometry. Show are the frequencies (upper panels) and geometric MFI (bottom panels) for CD4 **(A)** and CXCR4 **(B)** expression. CCR5 mRNA expression was quantified by RT-PCR **(C)**. Each symbol represents one donor (median ± interquartile range) for experiments performed with n=8 **(A-B)** an n=5 **(C)** participants. Friedman and uncorrected Dunn’s multiple comparison p-values are indicated on the graphs.

**Supplemental Figure 4: Metformin increases the expression of RORC2 and CCR6 on memory CD4^+^ T-cells HIV-infected *in vitro*.** (**A-G**) The HIV-infection *in vitro* was performed on memory CD4^+^ T-cells from HIV-uninfected donors (n=8), as described in Figure 3A. Shown are representative flow cytometry dot plots of intracellular RORC2 and surface CCR6 expression (**A)**; as well as statistical analysis of the frequency of RORC2^+^ T-cells **(B)** and the geometric MFIRORC2 expression **(C)**; the frequency of CCR6^+ T-^cells **(D)** and the geometric MFI of CCR6 expression **(E)**; as well as the frequency of Th17-like cells identified as cells with a CCR6^+^ RORC2^+^ phenotype **(F).** Shown is the kinetic IL-17A production quantified in the cell-culture supernatant by ELISA in one representative donor **(G, left panel)**, as well as statistical analysis of IL-17A production at day 9 posy-infection **(G, right panel)**. Finally, shown is the statistical analysis of the frequency of IL-17A^+^ T-cells **(H, left panel)** and the geometric MFI of intracellular IL-17A expression **(H, right panel)** at day 3 post-TCR triggering without HIV-1 infection for n=4 donors. Each symbol represents one donor; bars indicate the median ± interquartile range. Friedman and uncorrected Dunn’s multiple comparison p-values are indicated on the graphs.

**Supplemental Figure 5: Effect of metformin on anti-HIV Env binding to CD4^low^HIV-p24^+^ cells from ART-treated PWH.** The VOA was performed on memory CD4^+^ T-cells from ART-treated PWH, as described in Figure 1A. Shown is the gating strategy to identify by cytometry the anti-HIV-Env antibodies binding to CD4^low^HIV-p24^+^ T-cells in one representative donor **(A)**; dot plots of anti-HIV Env and HIV-p24 Abs binding on live cells of one HIV-uninfected donor, as a negative control for Abs specificity; and statistical analyses of the frequency of CD4l^ow^ T-cells co-expressing HIV-p24 intracellularly and HIV-Env on the surface of live cells **(C)**.

