## Supplemental Figures 1-5 for "Metformin Enhances Antibody-Mediated Recognition of HIV-Infected CD4^+^ T-Cells by Decreasing Viral Release"

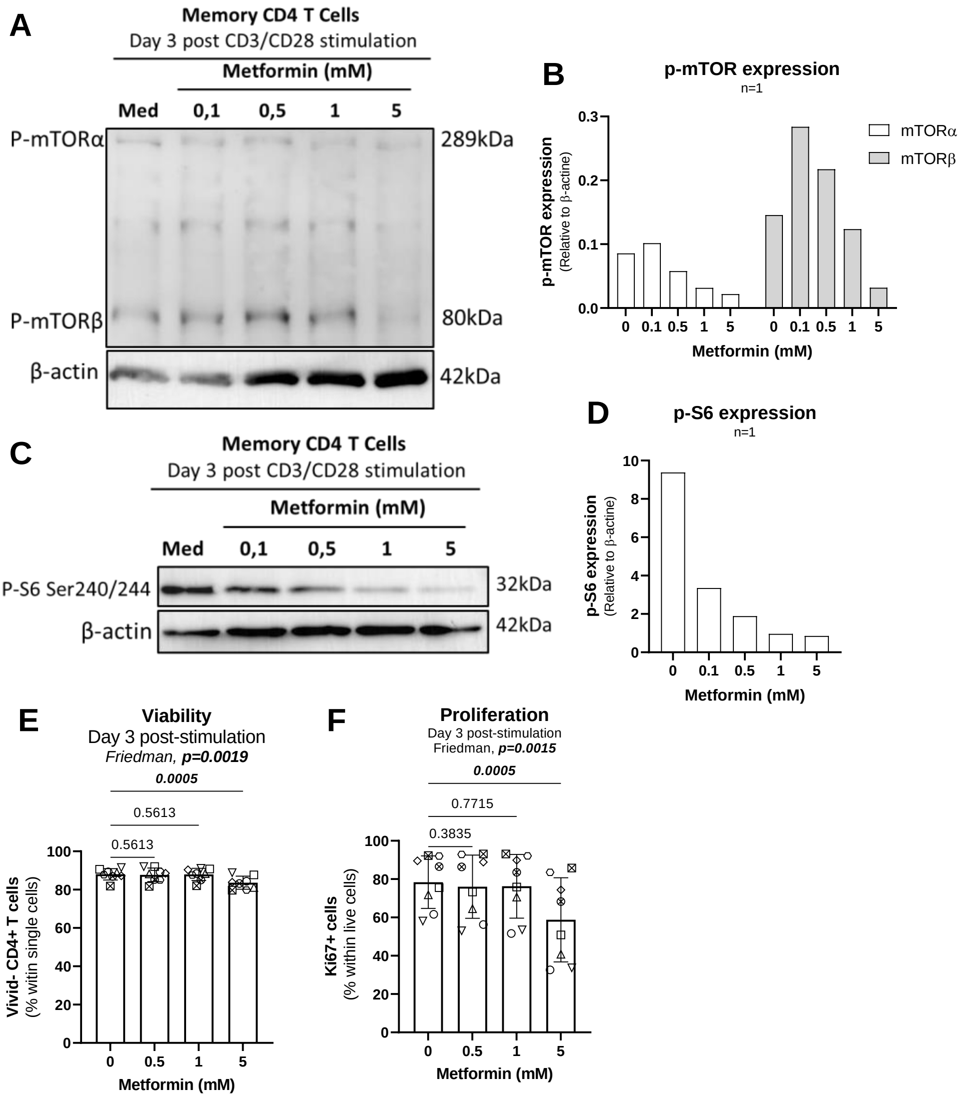


Fert *et al.,* Supplemental Figure 1 - Related to Figure 1

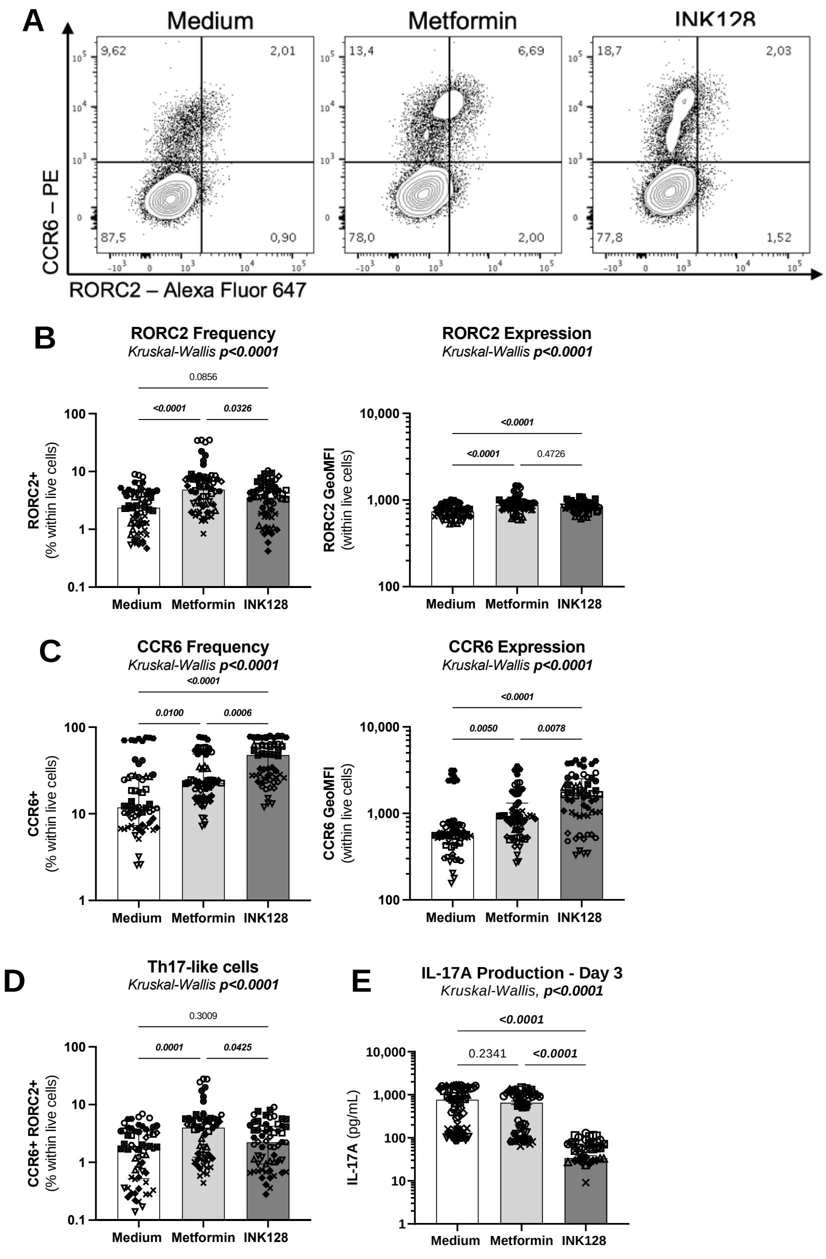


Fert *et al.,* Supplemental Figure 2 - Related to Figure 1

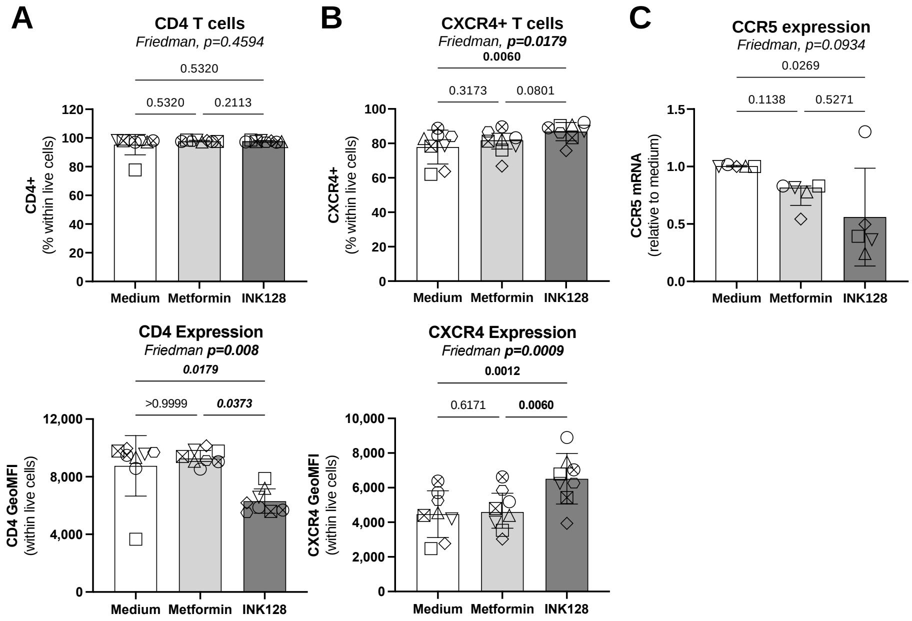


Fert *et al.,* Supplemental Figure 3 - Related to Figure 3

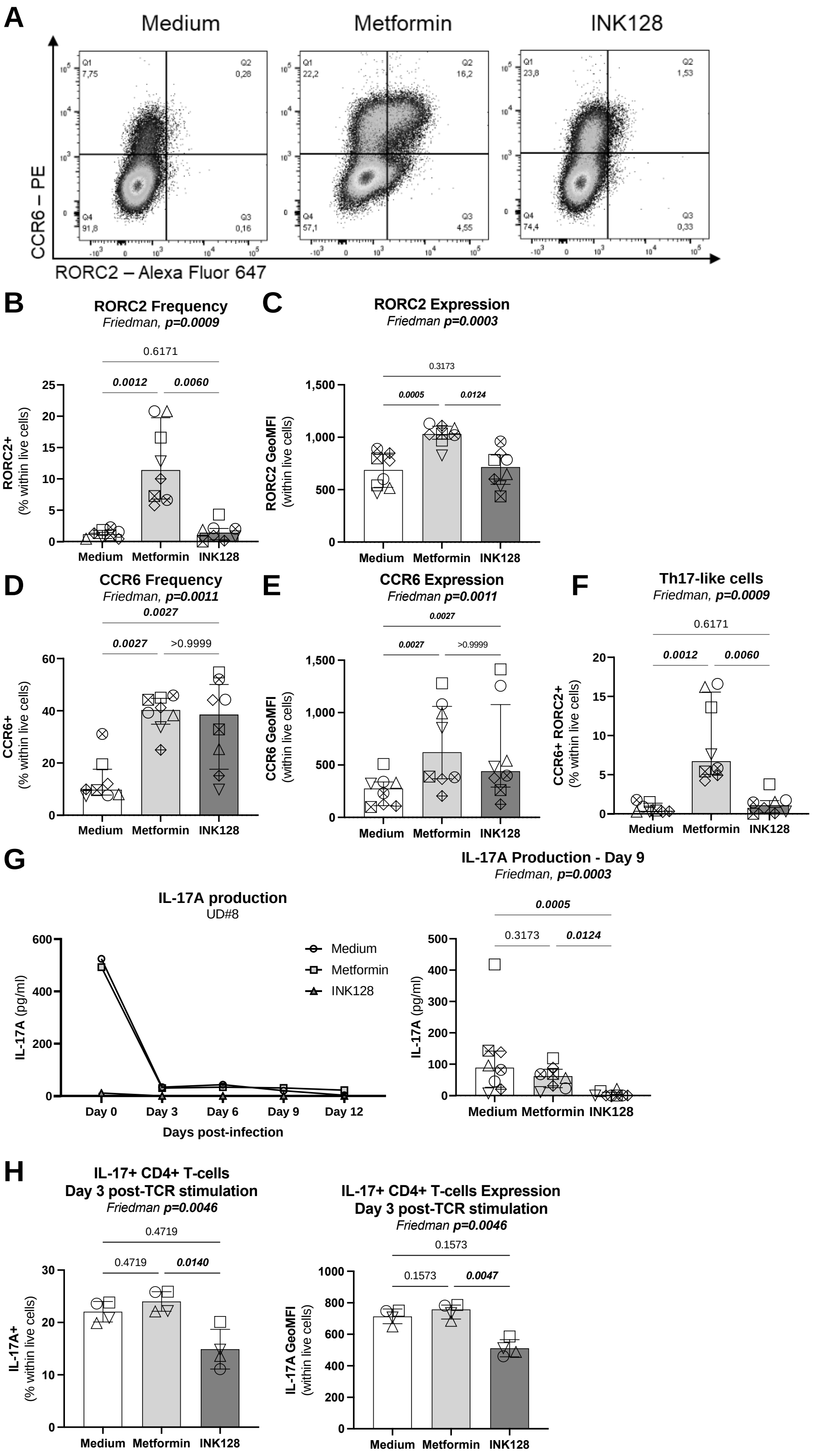


Fert *et al.,* Supplemental Figure 4 - Related to Figure 3

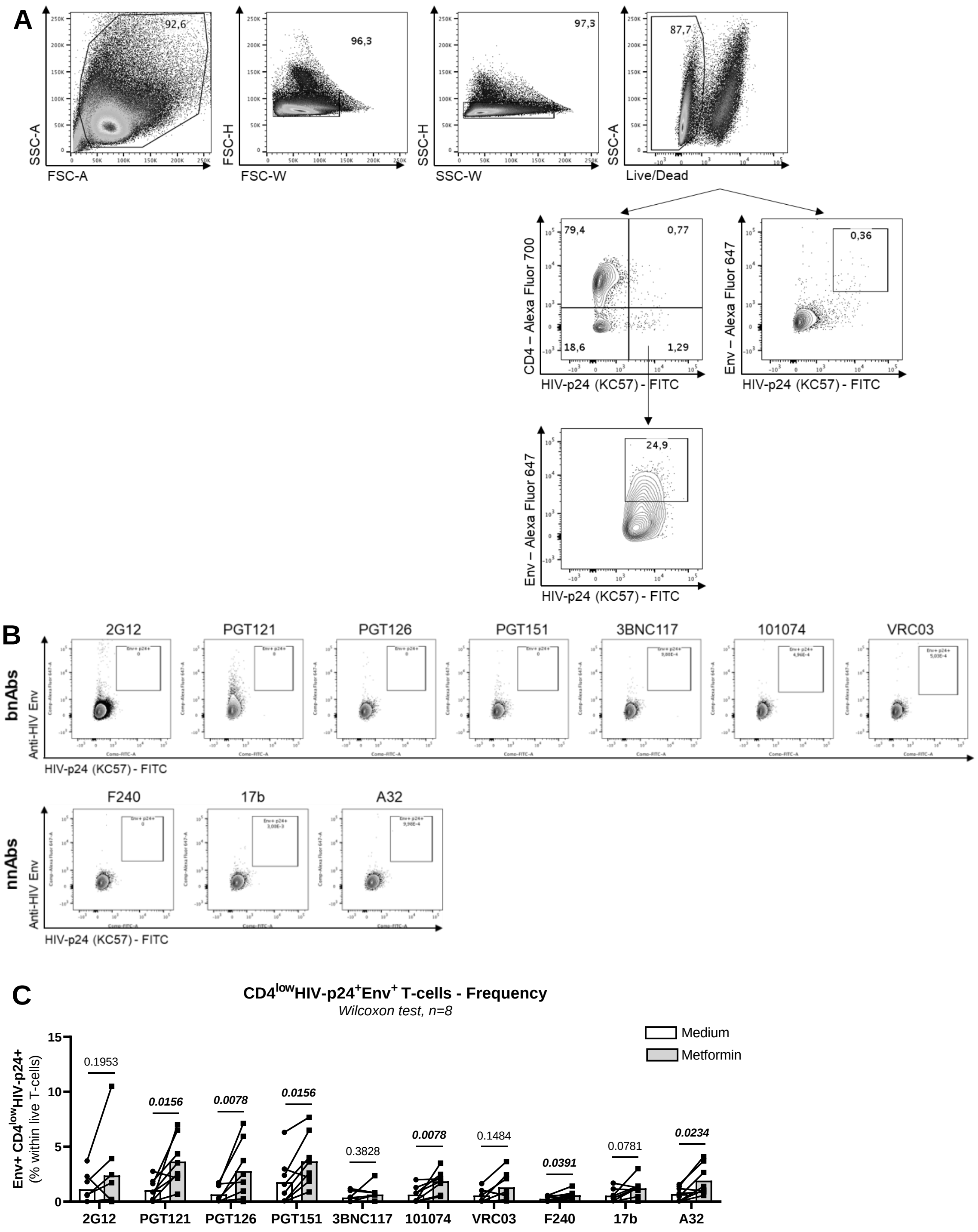


Fert *et al., S*upplemental Figure 5 - Related to Figure 7
