## Supplementary material for "Metformin Enhances Antibody-Mediated Recognition of HIV-Infected CD4^+^ T-Cells by Decreasing Viral Release": Key ressource Table

**Key Resources Table**

|  | SOURCE | IDENTIFIER |
| --- | --- | --- |
| **Antibodies** | | |
| Purified NA/LE Mouse Anti-Human CD3 (Clone UCHT1) | BD | Cat#555329; RRID: AB_395736 |
| Purified NA/LE Mouse Anti-Human CD28 (Clone CD28.2) | BD | Cat#555725; RRID: AB_396068 |
| Mouse anti-human CD3 Pacific Blue (Clone UCHT1) | BD | Cat#558117; RRID: AB_397038 |
| Mouse anti-human CD4 Alexa Fluor (Clone RPA-T4) | BD | Cat#561030; RRID: AB_10563215 |
| Mouse anti-human CD8 FITC (Clone 130-080-601) | Miltenyi | Cat#130-080-601; RRID: AB_244336 |
| Mouse anti-human CCR6 PE (Clone 11A9) | BD | Cat#559562; RRID: AB_397273 |
| Mouse anti-human CD45RA APC-H7 (Clone HI100) | BD | Cat#560674; RRID: AB_1727497 |
| Mouse anti-human CXCR4 APC (Clone 12G5) | eBiosciences | Cat#17-9999-42; RRID: AB_1724113 |
| Mouse anti-human CD317 (BST-2) BV421 (Clone Y129) | BD | Cat#566381; RRID: AB_2744363 |
| Hamster anti-human Bcl-2 FITC | BD | Cat#554234; RRID: AB_395319 |
| HIV-1 core (p24) antigen-FITC (Clone KC57) | Beckman Coulter | Cat#6604665 |
| HIV-1 core (p24) antigen-RD1 (Clone KC57) | Beckman Coulter | Cat#6604667; RRID: AB_1575989 |
| Mouse anti-human RORC2 Alex Fluor 647 (Clone Q21-559) | BD | Cat#563620; RRID: AB_2738324 |
| Mouse anti-human IL-17A PE (Clone eBio64DEC17) | eBiosciences | Cat#12-7179-42; RRID: AB_1724136 |
| Goat anti-Human IgG Secondary Antibody, Alexa Fluor 647 | Invitrogen | Cat#A-21445; RRID: AB_2535862 |
| LIVE/DEAD Fixable Aqua Dead Cell Stain Kit (405 nm excitation) | Invitrogen | Cat#L34957 |
| Anti-phospho-Ribosomal Protein S6 (Ser240/Ser244) | EMD Millipore | Cat#07 2113 |
| Phospho-mTOR (Ser2448) | Cell Signaling | Cat#2971; RRID: AB_330970 |
| Secondary antibody anti-Rabbit IgG HRP-linked | Cell Signaling | Cat#7074; RRID: AB_2099233 |
| Monoclonal Mouse Anti-β-Actin antibody | Millipore Sigma | Cat#A5441; RRID: AB_476744 |
| Goat anti-Mouse IgG (H+L) Secondary Antibody, HRP | Invitrogen | Cat#32430; RRID: AB_1185566 |
| **Virus Strains** | | |
| NL4.3BaL HIV plasmid | From Dr. Michel tremblay, Université Laval, Québec, Canada | N/A |
| pHEF Expressing Vesicular Stomatitis Virus (VSV-G) plasmid | NIH HIV Reagent Program, Division AIDS, NIAID (Contribution of Dr. Lung Ji Chang) | Cat#ARP-4693 |
| HIV-1 NL4-3 ΔEnv EGFP Reporter Vector | NIH HIV Reagent Program, Division of AIDS, NIAID (Contribution Dr. Haili Zhang, Dr. Yan Zhou and Dr. Robert Siliciano) | Cat#ARP11100 |
| **Biological Samples** | | |
| Leukaphereses of ART treated people living with HIV and HIV-uninfected people | Recruited at the Montreal Chest Institute, McGill University Health Centre and Centre Hospitalier de l’Université de Montréal with the help of Dr Jean-Pierre Routy’s group | N/A |
| **Chemicals and Cytokines** | | |
| rhIL-2 | R&D Systems | Cat#202-IL-050 |
| 1,1-Dimethylbiguanide, Hydrochloride (Metformin) | Santa Cruz | Cat#sc-202000 |
| INK128 (Cayman Chemical) | Cederlane | Cat#11811-1 |
| Saquinavir | NIH HIV Reagent Program, Division of AIDS, NIAID (Contribution DAIDS/NIAID) | Cat#ARP-4658 |
| Raltegravir | NIH HIV Reagent Program, Division of AIDS, NIAID (Contribution DAIDS/NIAID) | Cat#ARP-11680 |
| RIPA buffer | Cell Signaling | Cat#9806S |
| PhosSTOP (phosphatase inhibitors) | Milipore Sigma | Cat#4906845001 |
| cOmplete™, Mini, EDTA-free Protease Inhibitor Cocktail | Milipore Sigma | Cat#11836170001 |
| Clarity Max™ Western ECL Substrate | Bio-Rad | Cat# 1705062 |
| Re-Blot Plus Strong Solution | EMD Millipore | Cat#2504 |
| **Critical Commercial Assays** | | |
| EasySep Memory CD4+ T Cell Isolation Kit, human | StemCell Technology | Cat#19157 |
| Fixation/Permeabilization Solution Kit (Cytofix/Cytoperm) | BD | Cat#554714 |
| HIV-p24 ELISA | Homemade. Hybridome provided by Dr. Michel J. Tremblay (Bounou et al., J Virol., 2002) | |
| QuantiTect SYBR Green RT-PCR Kit | Qiagen | Cat #204245 |
| LC480 probe master mix | Roche | Cat# 4707494001 |
| SuperScript III One Step RT-PCR Taq polymerase | Invitrogen | Cat#12574-026 |
| PerfeCTa qPCR ToughMix, Low ROX de Quantabio | VWR | Cat# CA97065-970 |
| AllPrep DNA/RNA/miRNA Universal kit | Qiagen | Cat#80224 |
| Rneasy Plus Mini Kit | Qiagen | Cat#74136 |
| Wizard® Minipreps DNA Purification | Promega | Cat#A7100 |
| EndoFree Plasmid Maxi Kit | Qiagen | Cat#12362 |
| X-tremeGENE 9 kit | Roche | Cat#06 365 779 001 |
| DMTM Protein Assay | BioRad | Cat#500 0114 |
| **Culture media** | | |
| RPMI1640 | Gibco | Cat#11875-093 |
| Penicillin/Streptomycin | Gibco | Cat#15140-122 |
| Fetal Bovine Serum (FBS) | Wisent |  |
| **Experimental Models: Cell Lines** | | |
| ACH-2 cell line | NIH HIV Reagent Program, Division of AIDS, NIAID, NIH: contributed by Dr. Thomas Folks | Cat#ARP-349; RRID: CVCL_0138 |
| **Oligonucleotides** | | |
| See Supplemental Table 2 and 3 |  |  |
| **Software and Algorithms** | | |
| FlowJo version 10 | BD | https://www.flowjo.com/ |
| GraphPad Prism 9.0.1 | GraphPad | https://www.graphpad.com/ |
