## Supplemental Tables 1-3 for "Metformin Enhances Antibody-Mediated Recognition of HIV-Infected CD4^+^ T-Cells by Decreasing Viral Release"

**Supplemental Table 1**: Clinical parameters of HIV-uninfected study participants

| **Participants** | **Sex** | **Ethnicity** | **Age** |
| --- | --- | --- | --- |
| **HIV- #1** | N/A | N/A | N/A |
| **HIV- #2** | N/A | N/A | N/A |
| **HIV- #3** | N/A | N/A | N/A |
| **HIV- #4** | N/A | N/A | N/A |
| **HIV- #5** | M | Caucasian | 59 |
| **HIV- #6** | M | Caucasian | 59 |
| **HIV- #7** | M | N/A | 26 |
| **HIV- #8** | N/A | N/A | N/A |
| **HIV- #9** | N/A | N/A | N/A |
| **HIV- #10** | M | Caucasian | 60 |
| **HIV- #11** | M | Caucasian | 63 |
| **HIV- #12** | M | Caucasian | 44 |
| **HIV- #13** | M | Caucasian | 68 |
| **HIV- #14** | M | Caucasian | 32 |
| **HIV- #15** | M | Caucasian | 31 |

*HIV-, HIV-uninfected participants; N/A, information not available; M, male*

**Supplemental Table 2:** Oligonucleotides sequences of primers and probes used for early, late and integrated HIV-DNA real-time PCR quantification

|  |  | Types of PCR | Amplification | Sequence (5'→ 3') |
| --- | --- | --- | --- | --- |
| Primers | AA55 | RU5 | First | CGT CTA GAG ATT TTC CAC AC |
|  | M667 | RU5 | First | CTA ACT AGG GAA CCC ACT G |
|  | HCD3 OUT 5' | Integrated, RU5 & Gag | First | ACT GAC ATG GAA CAG GGG AAG |
|  | HCD3 OUT 3' | Proviral, RU5 & Gag | First | CCA GCT CTG AAG TAG GGA ACA TAT |
|  | Gag-R | Gag | First | AGC TCC CTG CTT GCC CAT A |
|  | Alu 1 | Integrated | First | TCC CAG CTA CTG GGG AGG CTG AGG |
|  | Alu 2 | INtegrated | First | GCC TCC CAA AGT GCT GGG ATT ACA G |
|  | LM667 | Integrated & Gag | First | ATG CCA CGT AAG CGA AAC TCT GGC TAA CTA GGG AAC CCA CTG |
|  | HIV Lambda T | Integrated & Gag | Second | ATG CCA CGT AAG CGA AAC T |
|  | AA55M | Integrated & Gag | Second | GCT AGA GAT TTT CCA CAC TGA CTA A |
|  | SK30 | RU5 | Second | GGT CTG AGG GAT CTC TAG |
|  | SK29 | RU5 | Second | ACT AGG GAA CCC ACT GCT |
|  | HCD3 IN 3' | Integrated, RU5 & Gag | Second | CCT CTC TTC AGC CAT TTA AGT A |
|  | HCD3 IN 5' | Integrated, RU5 & Gag | Second | GGC TAT CAT TCT TCT TCA AGG T |
| Probes | LTR-LC | Proviral & Total | Second | CAC TCA AGG CAA GCTT TAT TGA GGC |
|  | LTR-FL | Proviral & Total | Second | CAC AAC AGA CGG GCA CAC ACT ACT TGA |
|  | CD3-P1 | CD3 for normalisation in Integrated & Gag PCR | Second | GGC TGA AGG TTA GGG ATA CCA ATA TTC CTG TCT C |
|  | CD3-P2 | CD3 for normalisation in Integrated & Gag PCR | Second | CTA GTG ATG GGC TCT TCC CTT GAG CCC TTC |

**Supplemental Table 3:** Oligonucleotides sequences of primers and probes used for HIV transcription quantification

|  |  | Types of PCR | Amplification | Sequence (5'→3') |
| --- | --- | --- | --- | --- |
| Primers | ULF1 100 µM | Proviral & LTR-Gag RNA | First | ATG CCA CGT AAG CGA AAC TCT GGG TCT CTC TDG TTA G AC |
|  | UR1 100µM | Proviral & LTR-Gag RNA | First | CCA TCT CTC TCC TTC TAG C |
|  | HCD3 OUT 5' | CD3 for normalisation in Proviral PCR | First | ACT GAC ATG GAA CAG GGG AAG |
|  | HCD3 OUT 3' | CD3 for normalisation in Proviral PCR | First | CCA GCT CTG AAG TAG GGA ACA TAT |
|  | Alu 1 | Proviral | First | TCC CAG CTA CTG GGG AGG CTG AGG |
|  | Alu 2 | Proviral | First | GCC TCC CAA AGT GCT GGG ATT ACA G |
|  | HIV Lambda T | Proviral & LTR-Gag RNA | Second | ATG CCA CGT AAG CGA AAC T |
|  | UR2 | Proviral & LTR-Gag RNA | Second | CTG AGG GAT CTC TAG TTA CC |
|  | HCD3 IN 3' | CD3 for normalisation in Proviral PCR | Second | CCT CTC TTC AGC CAT TTA AGT A |
|  | HCD3 IN 5' | CD3 for normalisation in Proviral PCR | Second | GGC TAT CAT TCT TCT TCA AGG T |
| Probes | UHIV Famzen | Proviral & LTR-Gag RNA | Second | 56-FAM/CA CTC AAG G/ZEN/C AAG CTT TAT TGA GGC /3IABkFQ/ |
|  | CD3 Famzen | CD3 for normalisation in Proviral PCR | Second | /56-FAM/AG CAG AGA A/ZEN/C AGT TAA GAG CCT CCA T/3IABkFQ/ |
